# GOT1 Inhibition Primes Pancreatic Cancer for Ferroptosis through the Autophagic Release of Labile Iron

**DOI:** 10.1101/2020.02.28.970228

**Authors:** Daniel M. Kremer, Barbara S. Nelson, Lin Lin, Emily L. Yarosz, Christopher J. Halbrook, Samuel A. Kerk, Peter Sajjakulnukit, Amy Myers, Galloway Thurston, Sean W. Hou, Eileen S. Carpenter, Anthony C. Andren, Zeribe C. Nwosu, Nicholas Cusmano, Stephanie Wisner, Johanna Ramos, Tina Gao, Stephen A. Sastra, Carmine F. Palermo, Michael A. Badgley, Li Zhang, John M. Asara, Marina Pasca di Magliano, Yatrik M. Shah, Howard C. Crawford, Kenneth P. Olive, Costas A. Lyssiotis

## Abstract

Pancreatic ductal adenocarcinoma (PDA) is one of the deadliest solid malignancies, with a 5-year survival rate at ten percent. PDA have unique metabolic adaptations in response to cell-intrinsic and environmental stressors, and identifying new strategies to target these adaptions is an area of active research. We previously described a dependency on a cytosolic aspartate aminotransaminase (GOT1)-dependent pathway for NADPH generation. Here, we sought to identify metabolic dependencies induced by GOT1 inhibition that could be exploited to selectively kill PDA. Using pharmacological methods, we identified cysteine, glutathione, and lipid antioxidant function as metabolic vulnerabilities following GOT1 withdrawal. Targeting any of these pathways was synthetic lethal in GOT1 knockdown cells and triggered ferroptosis, an oxidative, non-apoptotic, iron-dependent form of cell death. Mechanistically, GOT1 inhibition promoted the activation of autophagy in response to metabolic stress. This enhanced the availability of labile iron through ferritinophagy, the autolysosome-mediated degradation of ferritin. In sum, our study identifies a novel biochemical connection between GOT1, iron regulation, and ferroptosis, and suggests the rewired malate-aspartate shuttle plays a role in protecting PDA from severe oxidative challenge.

**Highlights:**

- PDA exhibit varying dependence on GOT1 for *in vitro* and *in vivo* growth.
- Exogenous cystine, glutathione synthesis, and lipid antioxidant fidelity are essential under GOT1 suppression.
- GOT1 inhibition sensitizes pancreatic cancer cell lines to ferroptosis.
- GOT1 inhibition represses anabolic metabolism and promotes the release of iron through autophagy.

## Introduction

Pancreatic ductal adenocarcinoma (PDA) is a notoriously lethal disease. This stems from late-stage diagnosis and the lack of effective therapies^1^. A defining feature of PDA is the extensive fibroinflammatory reaction that not only regulates cancer initiation, progression, and maintenance, but also promotes its therapeutic resilience^2,3,4^. Vascular collapse, impaired perfusion, and hypoxia accompany this desmoplastic reaction to produce a nutrient-deprived and harsh tumor microenvironment^3,5^. PDA cells reprogram their nutrient acquisition and metabolism to support survival and growth under these conditions^6,7^.

Our previous work demonstrated that PDA rewire the malate-aspartate shuttle to generate reduced nicotinamide adenine dinucleotide phosphate (NADPH), a major currency for biosynthesis and redox balance **(Figure 1A)**^8^. The malate-aspartate shuttle canonically functions to transfer reducing equivalents in the form of NADH from the cytosol into the mitochondria to facilitate oxidative phosphorylation (OxPHOS). In PDA, we found the mitochondrial aspartate aminotransaminase (GOT2) is the primary anaplerotic source for alpha-ketoglutarate (αKG) and generates aspartate. Aspartate is then transferred to the cytosol and transaminated to produce oxaloacetate (OAA) by the cytosolic aspartate aminotransaminase (GOT1). OAA is reduced to malate by cytosolic Malate Dehydrogenase (MDH1) and is then oxidized by Malic Enzyme 1 (ME1) to generate NADPH, which is utilized to support redox balance and proliferation in PDA^8^. Furthermore, we demonstrated that this non-canonical pathway was orchestrated by mutant KRAS, the signature oncogenic driver of PDA. Specifically, mutant Kras led to the transcriptional upregulation of GOT1 and the concurrent repression of GLUD1, which drives an alternative metabolic pathway. Indeed, a higher expression of *GOT1* to *GLUD1* is associated with worse patient outcome^9^. Thus, in an effort to target this rewired metabolic pathway, we have placed our focus on GOT1. And, notably, we and others recently identified drug scaffolds that may serve as leads in the development of a clinical grade GOT1 inhibitor ^10,11,12,13^.

**Figure 1.**
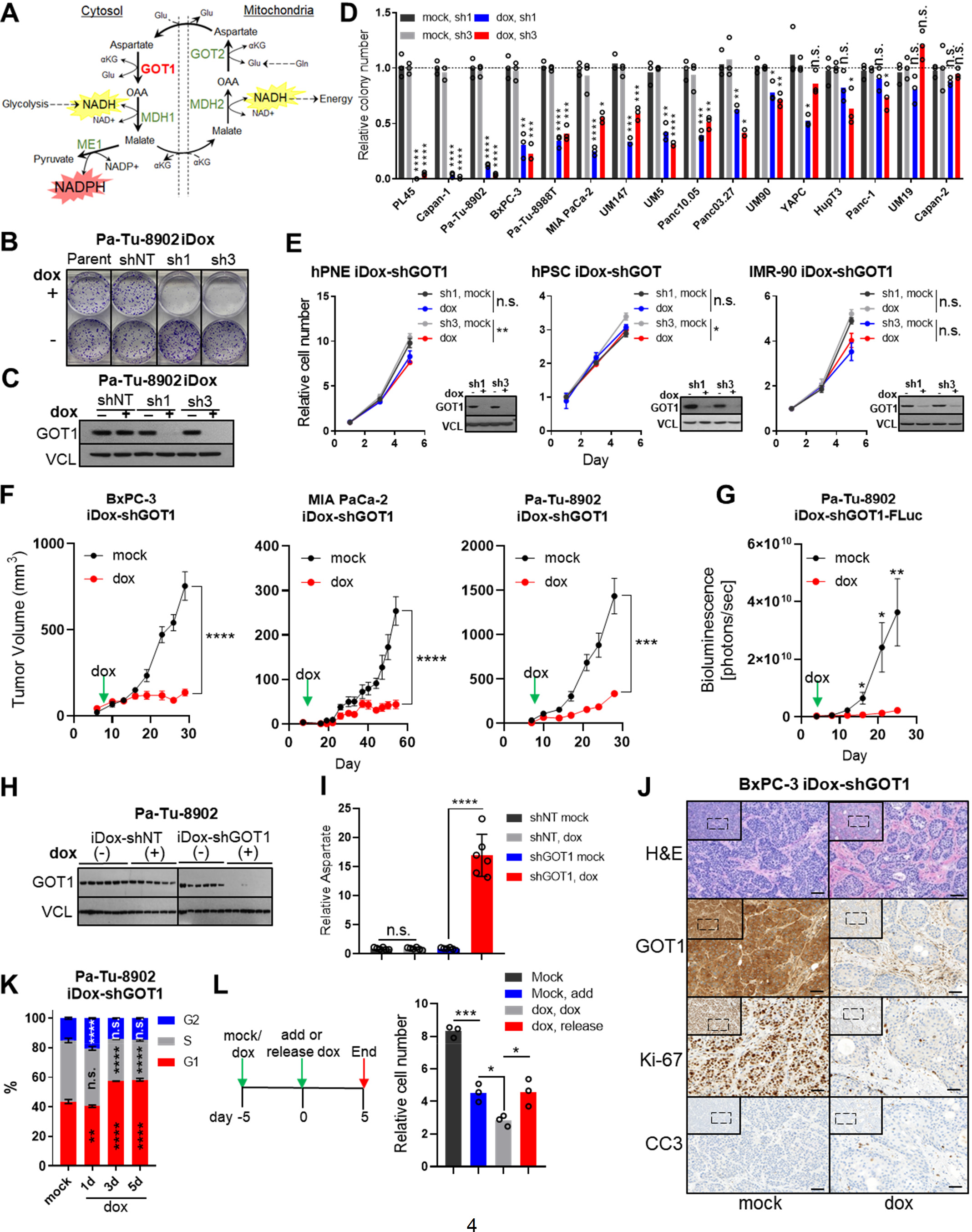
GOT1 dependence is a feature of some PDA cell lines, matched by G1 cell cycle arrest and inhibition of proliferation. **A**) Model of the rewired malate-aspartate shuttle in PDA. **B-C**) Representative colony formation assay (**B**) and immunoblot analysis (**C**) of Pa-Tu-8902 cells stably expressing iDox-shRNA constructs following 10 days GOT1 knockdown. shRNAs target the coding region of GOT1 (sh1), or the 3’UTR region of GOT1 (sh3). Parental (parent) and scramble (shNT) conditions are also displayed (n=3). Vinculin (VCL) was used as a loading control. **D**) Relative colony number across a panel of PDA cell lines in response to GOT1 knockdown by sh1 (black/blue) or sh 3 (grey/red). Assays were run 10-15 days (n=3), n.s. denotes non-significant. **E**) Relative cell number of immortalized, non-transformed, cell lines after 1, 3, or 5 days, normalized to day 1 (n=3). **F**) Growth of subcutaneous xenograft tumors from 3 PDA cell lines stably expressing iDox-shGOT1 or shNT. Dox or mock conditions were administered 7 days following implantation. Treatment with dox (red) or vehicle (black) (BxPC-3 n= 8, MIA PaCa-2 n=6, Pa-Tu-8902 n=6 per arm). **G**) Orthotopic xenograft tumor growth from Pa-Tu-8902 iDox-shGOT1 stable cell lines coexpressing firefly luciferase (FLuc) n=5 and n=6 mice were used for vehicle and dox cohorts respectively. **H-I**) Immunoblots for GOT1 (**H**) and relative aspartate levels (**I**) measured by liquid-chromatography tandem mass spectrometry, normalized to-dox (n=5). **J**) Histology of BxPC-3 iDox-shGOT1 subcutaneous xenograft tumors from vehicle-or dox-treated mice. H&E, Hematoxylin and Eosin, CC3, cleaved caspase 3. Scale bars represent 50μm. **K**) Cell cycle distribution of Pa-Tu-8902 iDox-GOT1 sh1 upon 1,3, or 5 days of dox treatment. Significance values are in relation to iDox-shGOT1 mock (n=3). **L**) Proliferation kinetics following GOT1 knockdown. Cells were untreated (black), dox was added to untreated cells (blue), pretreated with dox and chronically exposed to dox (grey), or released from dox pretreated cells (red). Relative cell number at day 5 normalized to day 1 is displayed, (n=3). Error bars represent mean ± SD. Two-tailed unpaired T-test or 1-way ANOVA: Non-significant P > 0.05 (n.s. or # as noted), P ≤ 0.05 (*), ≤ 0.01 (**), ≤ 0.001 (***), ≤ 0.0001 (****). See also **Figure S1**.

Herein, we present a detailed analysis of GOT1 dependence across a large panel of PDA cell lines and specimens. We demonstrate that GOT1 sensitivity varies among the cultures in this panel, is dispensable in non-transformed human lines, and that GOT1 inhibition stunted growth in tumor xenograft models. In GOT1 dependent contexts, GOT1 inhibition blocked progression through the cell cycle, leading to cytostasis. Thus, we then sought to characterize metabolic dependencies following GOT1 withdrawal that could be exploited to selectively kill PDA^14,15^. Examination of a targeted library of metabolic inhibitors in GOT1 knockdown cells led to the discovery that that exogenous cystine was essential for viability following chronic GOT1 suppression. Cystine is used for reduced glutathione (GSH) biosynthesis, which mediates protection against lipid oxidation. GOT1 knockdown in combination with inhibitors of glutathione synthesis or lipid antioxidant machinery led to cell death. We characterized this as ferroptosis: an oxidative, non-apoptotic, and iron-dependent form of cell death. We then determined that GOT1 withdrawal promoted a catabolic cell state that resulted in decreased OxPHOS and the activation of autophagy and ferritinophagy^16^. Ferritinophagy primed GOT1 knockdown cells for ferroptosis by increasing labile iron pools. Together with the modulation of NADPH and GSH levels, our study demonstrates that GOT1 inhibition promotes ferroptosis sensitivity by promoting labile iron and illustrates how the rewired malate-aspartate shuttle participates in the maintenance of both antioxidant and energetic balance.

## Results

### GOT1 dependent PDAs require GOT1 for cell cycle progression and proliferation

To examine GOT1 dependence in a large panel of PDA cell lines and primary specimens with temporal control, we developed doxycycline (dox)-inducible short hairpin (sh)RNA reagents (iDox-sh) that target the coding and 3’UTR regions of GOT1 (sh1 and sh3), or scramble (shNT). shRNA activity was examined phenotypically by assessing colony formation and protein levels following dox treatment **(Figures 1B,C and S1A,B)**. We also measured aspartate levels as a biochemical readout for GOT1 inhibition **(Figure S1C)**. We then used these iDox-shRNA constructs to assess GOT1 sensitivity across a large panel of PDA lines and primary specimens (indicated with the UM# designation)^17^ **(Figure 1D and S1D)**. GOT1 knockdown significantly impaired colony formation in 12 of 18 cell lines in the panel **(Figures 1D and S1A,B)**.

Indeed, in PL45, Capan-1, and Pa-Tu-8902 colony formation was severely diminished following GOT1 knockdown with both hairpins, and this was independent of dox-effects **(Figure S1D)**. Further, the response to GOT1 knockdown did not correlate with common mutations associated with PDA **(Figure S1E)** or expression of malate-aspartate shuttle enzymes **(Figure S1F)**. To test the specificity of GOT1 against PDA, we extended our cell panel to non-transformed human lines. We found human pancreatic stellate cells (hPSC), human lung fibroblasts (IMR-90), and human non-transformed pancreatic exocrine cells (hPNE) were minimally affected upon GOT1 knockdown, in agreement with previous results, suggesting that this pathway may be dispensable in non-transformed cells **(Figures 1E and S1G)**^8,12^. Together, these data demonstrate many PDA cell lines require GOT1 for growth while non-transformed cell lines do not, highlighting a potential therapeutic window.

To determine the relevance of GOT1 *in vivo*, we examined the effect of GOT1 inhibition on established PDA tumors. PDA cells were implanted subcutaneously into the flanks or orthotopically into the pancreas of immunocompromised mice and allowed to establish for 7 days prior to GOT1 inhibition. GOT1 sensitive cell lines exhibited profound growth inhibition upon induction of GOT1 knockdown with dox (Figures 1F,G), results that were consistent with previous studies that employed constitutive shGOT1^8^. Parallel studies with shNT tumors indicated that the effect was independent of dox exposure **(Figure S1H)**. GOT1 knockdown was demonstrated by immunoblot analysis on homogenized tumor tissue **(Figures 1H and S1I)** and biochemically via the induction of aspartate **(Figure 1I)**. Contrasting our *in vitro* studies in which GOT1 was knocked down for five days **(Figure S1C)**, the changes in aspartate abundance described in this experiment reflect the whole tumor metabolome after thirty days of dox treatment. While immunostaining for GOT1 indicate potent knockdown in the PDA cell compartment **(Figures 1J)**, it is unknown how long-term suppression of GOT1 would influence non-cell autonomous metabolism in this xenograft study. Tumor growth suppression was confirmed at the molecular level by a decrease in Ki-67, a marker for proliferation. GOT1 knockdown tumors exhibited minimal staining for cleaved caspase 3 (CC3), a marker for apoptosis. Thus, these proliferative defects were independent of apoptosis, indicating GOT1 inhibits tumor proliferation, rather than, inducing cell death **(Figure 1J)**.

To test the hypothesis that GOT1 inhibition is cytostatic, we examined the effect of GOT1 knockdown on cell cycle progression. Knockdown led to a higher distribution of cells in G1 phase versus the S and G2 phases, indicating that the majority of cells are in G1 cell cycle arrest following GOT1 inhibition **(Figures 1K and S1J)**. Moreover, the effect of GOT1 knockdown was reversible, as cells regained proliferative capacity upon removal of genetic inhibition **(Figures 1L and S1K)**. Overall, PDA display a spectrum of sensitivity to GOT1 where GOT1 inhibition is cytostatic.

### Limiting exogenous cystine potentiates GOT1 inhibition and elicits a cytotoxic effect

Because GOT1 inhibition is cytostatic, we sought to identify metabolic dependencies induced by knockdown that could be targeted to selectively kill PDA^14,15^. Our previous work indicated that inhibiting glutamine metabolism enhances sensitivity to reactive oxygen species (ROS)^8,9,18^. Thus, we assessed the sensitivity of PDA cells to a panel of chemotherapeutic agents and inhibitors of antioxidant pathways after five days of GOT1 knockdown and three days of drug treatment **(Figures 2A,B)**.

**Figure 2.**
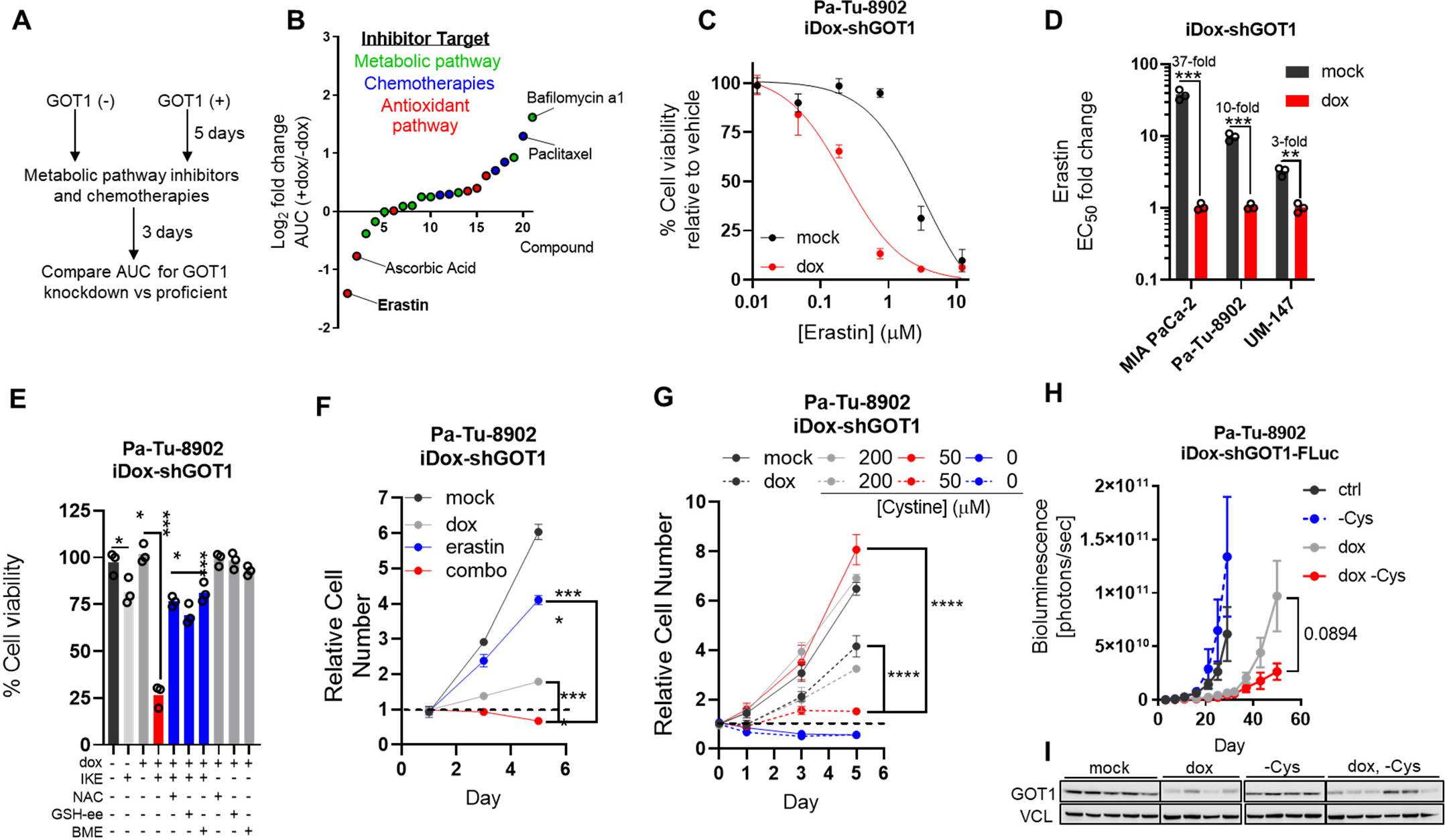
Limiting exogenous cystine causes a cytotoxic response to GOT1 inhibition. **A**) Screening strategy to identify metabolic dependencies induced by GOT1 knockdown. Pa-Tu-8902 iDox-shGOT1 cells were treated with vehicle or dox for 5 days followed by addition of the compounds for 72 h. **B**) Log_2_ fold change in area under the curve (AUC) from cell viability dose response curves for each compound in the library. AUC from GOT1 knockdown conditions (dox) are relative to (mock) conditions. Sensitivity ranking (Rank) spans 1-21, signifying the most sensitive (1) to least sensitive (21) compounds in response to knockdown (n=3). **C**) Dose-response curves for Pa-Tu-8902 iDox-shGOT1 cells following 5 days of GOT1 knockdown and erastin for 24 hours (n=3). **D**) EC_50_ values relative to +dox for 3 iDox-shGOT1 PDA cell lines following 24 hours of erastin treatment (n=3). **E**) Relative Pa-Tu-8902 iDox-shGOT1 cell numbers following 5 days of GOT1 knockdown with the indicated media conditions for 24 hours. 750nM of Erastin was administered on day 1 and conditions are normalized to day 1 (n=3). **F**) Cell viability of Pa-Tu-8902 iDox-shGOT1 after 5 days of GOT1 knockdown then 24 hours of 750nM IKE combined with the indicated conditions. 250μM of N-acetyl-cysteine (NAC), 250μM GSH-ethyl ester (GSH-EE), and 50μM of beta-mercaptoethanol (BME) were used (n=3). **G**) Relative Pa-Tu-8902 iDox-shGOT1 cell numbers following 5 days of GOT1 knockdown and the indicated media conditions (n=3). 200μM of cystine is a supraphysiological concentration, while 50μM is tumor relevant. **H-I**) Orthotopic xenograft tumor growth from Pa-Tu-8902 iDox-shGOT1 stable cell lines co-expressing firefly luciferase (FLuc) treated with vehicle (black, n=6), dox containing food (red, n=6), cysteine-free diet (grey, n=5), or dox containing, cysteine-free food (blue, n=6). GOT1 immunoblot (**I**) was taken from endpoint tumors. Error bars represent mean ± SD in (**B-G**) or mean ± SEM in (**H**). Two-tailed unpaired T-test or 1-way ANOVA: Non-significant *P* > 0.05 (n.s. or # as noted), *P* ≤ 0.05 (*), ≤ 0.01 (**), ≤ 0.001 (***), ≤ 0.0001 (****). See also **Figure S2**.

GOT1 inhibition was protective when combined with some inhibitors demonstrated by increased area under the curve values (AUC), a measure of drug sensitivity **(Figures 2B and S2A)**. Three of the five top desensitizing agents were chemotherapies, in agreement with previous observations^18^. Since these chemotherapies work through disrupting DNA replication of rapidly dividing cells, we speculate that the decreased sensitivity may occur due to the GOT1 growth-suppressive phenotype **(Figures 1J,K)**. By contrast, GOT1 knockdown sensitized PDA cells to erastin **(Figure 2B)**, and the effect was stronger after 24 hours of treatment **(Figure 2C)**. Erastin is an inhibitor of xCT^19,20^, a component of the system x_c_^-^ cystine/glutamate antiporter which transports cystine into cells in exchange for glutamate. Cystine, the oxidized dimer of cysteine, is reduced to cysteine upon entering the cell where it can contribute to the synthesis of GSH and proteins, among numerous other biochemical fates.

We then tested the combination of GOT1 knockdown and erastin in a small panel of additional PDA lines **(Figure 2D)** and observed that GOT1 inhibition increased sensitivity to Erastin by greater than 10-fold in some cases **(Figure 2C and S2B)**. GOT1 knockdown also sensitized PDA to imidazole ketone erastin (IKE), an erastin analog with increased potency^21^ **(Figure S2D)**. Furthermore, treating cells with a nano-molar dose of erastin or IKE combined with GOT1 knockdown was cytotoxic, while single treatment arms were cytostatic **(Figures 2E and S2E)**, and these erastin doses were sufficient to reduce GSH levels after 6 hours **(Figure S2F)**. Because erastin and IKE disrupt the import of cystine into the cell, we sought to inhibit cytotoxicity by co-treating with exogenous cysteine sources or GSH. Indeed, supplementation with N-acetyl cysteine (NAC), β-mercaptoethanol (BME), or cell permeable GSH ethyl-ester (GSH-EE) prevented cell death **(Figures 2F and S2G)**, consistent with the concept that cystine import through system xc^-^ is essential to maintain GSH levels^22^.

Previous studies have found cystine levels to be limiting in the PDA microenvironment. Cysteine was shown to be the second most depleted amino acid in pancreatic tumors relative to adjacent healthy pancreatic tissue^23^ and the levels of cystine in PDA tumor interstitial fluid (~50μM) which is 2-fold lower than in plasma taken from the same tumor-bearing mice^24^. Based on these observations, we sought to test the effect of GOT1 knockdown under physiological concentrations of cystine. Culturing cells in tumor-relevant cystine concentrations was growth inhibitory, while cystine deprivation was cytotoxic **(Figures 2G and S2H)**. Moreover, cell viability decreased in a dose-dependent manner after, and the response was potentiated by GOT1 knockdown at lower cystine concentrations **(Figure S2I)**, in agreement with our pharmacological studies. These results suggest PDA require exogenous cystine for growth and cell viability following GOT1 inhibition.

To test this concept *in vivo*, we engrafted Pa-Tu-8902 iDox-shGOT1 cells engineered to express firefly luciferase (FLuc) into the pancreas, as in **Figure 1G**. Tumors were allowed to establish for 7 days, and treatment arms were initiated by providing dox-containing food formulated with or without the non-essential amino acid cysteine. While tumors in the animals fed a cysteine-free diet grew at comparable rates to tumors in animals fed a control diet, dox treated tumors grew substantially slower **(Figures 2H,I and S2J)**. Mice fed with a cysteine-free diet had lower cysteine in tumors compared to the control diet **(Figure S2K)**, indicating dietary inputs can influence tumor metabolism. By contrast, cysteine was not significantly altered in tumors comparing dox-single and double treatment arms **(Figure S2J)**. The difference in tumor growth or tumor burden for animals on the cysteine-free diet were markedly smaller at end point, though this did not reach statistical significance **(Figures 2H,I)**.

Overall, these data indicate that PDA cultures require exogenous cystine following GOT1 inhibition. We also demonstrate that this mechanism is operative *in vivo*, through PDA tumors may acquire cysteine through alternative mechanisms when challenged by chronic cysteine deprivation.

### Inhibiting GSH biosynthesis potentiates the growth inhibitory effects of GOT1 knockdown

Our data indicate that PDA are heavily reliant on exogenous cystine following GOT1 inhibition. Previous work from our group has demonstrated that one of the downstream effects of GOT1 is reducing oxidized glutathione (GSSG)^8,18^. Thus, we hypothesized that cysteine acquisition was upregulated for GSH biosynthesis to compensate for the loss of NADPH-mediated GSH salvage through the GOT1 pathway **(Figure 3A)**. To test this hypothesis, we inhibited *de novo* GSH biosynthesis in combination with GOT1 knockdown. The rate-limiting step in GSH synthesis is catalyzed by glutamate-cysteine ligase (GCL), which forms gamma glutamyl-cysteine through the condensation of glutamate and cysteine^25^.

**Figure 3.**
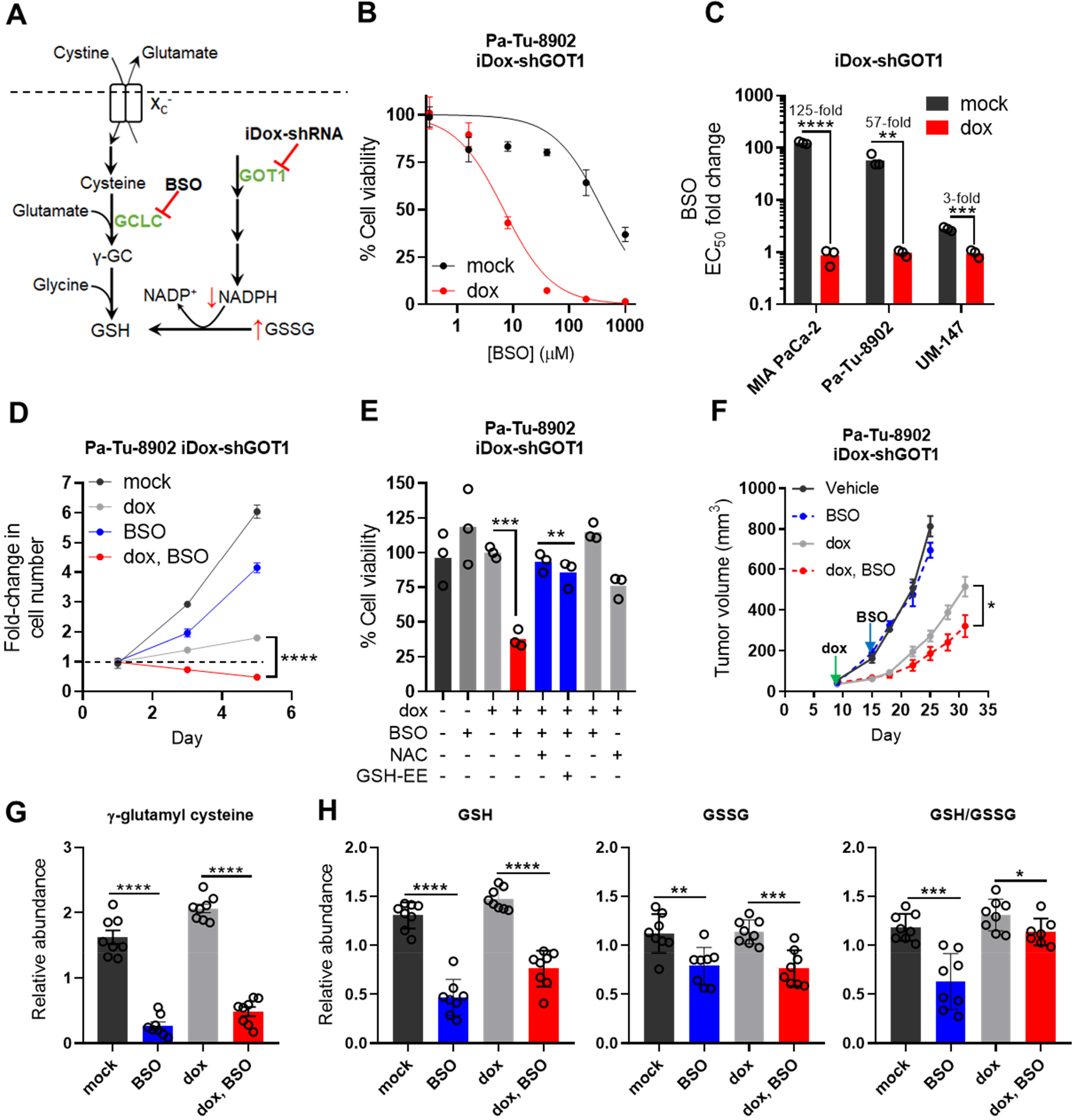
Inhibiting GSH biosynthesis produces a cytotoxic response upon GOT1 knockdown. **A**) Schematic integrating the GSH biosynthesis and GOT1 pathways. **B**) Percent cell viability dose-response curves upon 72 hours of BSO treatment following 5 days of GOT1 knockdown (n=3). **C**) EC_50_ values relative to +dox for 3 iDox-shGOT1 PDA cell lines following 72 hours of BSO treatment (n=3). **D**) Fold change Pa-Tu-8902 iDox-shGOT1 cell numbers following 5 days of GOT1 knockdown and treatment with the indicated conditions. 40 μM of BSO was administered on day 1. Cell numbers are normalized to day 1 for each condition (n=3). **E**) Percent cell viability following 72 hours of 40uM BSO or co-treatment with 0.5mM N-acetyl cysteine (NAC) or 0.5mM GSH-Ethyl Ester (GSH-EE) following 5 days of GOT1 knockdown (n=3). **F**) Subcutaneous xenograft growth of Pa-Tu-8902 iDox-shGOT1 cells treated with vehicle (black), 20 mg/kg BSO via drinking water (grey), doxycycline administered in the food (red), or the combination (blue). **G**) Relative abundance of gamma-glutamyl cysteine (yGC) from tumors in (**F**) (n=8). **H**) Relative abundances of GSH, GSSG, and the GSH/GSSG ratio from tumors in (**F**) (n=8). Error bars represent mean ± SD. Two-tailed unpaired T-test or 1-way ANOVA: Non-significant *P* > 0.05 (n.s. or # as noted), *P* ≤ 0.05 (*), ≤ 0.01 (**), ≤ 0.001 (***), ≤0.0001 (****). See also **Figure S3**.

GCL is a holoenzyme which consists of a catalytic subunit (GCLC) and a modifier subunit (GCLM), and the GCLC subunit is targeted by the inhibitor buthionine sulfoximine (BSO), resulting in decreased GSH production^26^ **(Figure 3A)**. We observed that GOT1 knockdown enhanced sensitivity to BSO after 24 hours of drug treatment, contrasting an absent single agent response **(Figure S3A)**. Exposure to BSO for 72 hours further augmented the sensitizing effect **(Figures 3B and S3B,C)**. In some cases, the change in EC50 was nearly 100-fold **(Figures 3C and S3D),** indicating potent sensitization.

Recent studies have suggested that a majority of cancer cell lines survive upon 72 hours of BSO treatment despite potent inhibition of GSH levels at this time point. Moreover, these studies have demonstrated that chronic BSO treatment, up to 9 days, is required in induce cell death or stasis^27^. In our models, 6 hours of BSO treatment was sufficient to diminish GSH levels **(Figure S3E)**, in line with previous kinetic data^27^. By contrast, co-treatment of GOT1 knockdown with BSO was cytotoxic at 72 hours, while BSO alone was cytostatic **(Figures 3D and S3F)**. 120 hours of treatment potentiated cell death in the GOT1 knockdown condition, whereas cells regained proliferative capacity under the BSO single treatment condition **(Figure 3D and S3F)**. Cell death could be prevented by supplementing exogenous GSH-EE or NAC **(Figures 3E and S3G)**, suggesting that cell death is due to perturbing GSH levels and redox balance.

Targeting GSH production with BSO is tolerated in patients, and BSO has been used in combination with chemotherapy in multiple phase 1 clinical trials, (NCT00005835 and NCT00002730)^28,29^. Thus, we sought to determine whether this combination shows efficacy *in vivo*. We examined the effect of GOT1 and BSO in established xenograft tumors. Mice were engrafted with Pa-Tu-8902 iDox-shGOT1 cells and given dox via chow after 7 days. BSO was administered via drinking water on day 14. While no tumor regressions were observed, the combination of GOT1 and BSO significantly slowed tumor progression compared with single treatment arms **(Figure 3F)** and led to complete stasis in one instance **(Figure S3H)**. Knockdown was confirmed immunoblot analysis **(Figure S3I)** and by LC-MS measurements of aspartate **(Figure S3J)** and on whole-tumor samples.

We then measured glutathione species in tumor metabolite fractions to demonstrate the pharmacodynamics of BSO. We found BSO to significantly reduce levels of gamma glutamyl-cysteine, a product of GCL, which is directly inhibited by BSO^25^ **(Figure 3G)**. Concomitantly, we observed a significant reduction in GSH, GSSG, and the GSH/GSSG ratio upon BSO treatment **(Figure 3H)**, demonstrating BSO has on-target activity in established tumors, and the tumors are under redox stress. Together, our data reveal that PDA require glutathione synthesis under GOT1 deficient conditions.

### GOT1 suppression sensitizes PDA to ferroptosis

Previous work has demonstrated that some cell types are sensitive to erastin and BSO as single agents, and that these drugs can kill cells by depleting GSH. The proximal effects of GSH depletion are mediated through loss of GPX4 activity, which utilizes GSH as a co-factor to detoxify lipid peroxides **(Figure 4A)**. This can lead to the lethal accumulation of lipid peroxides, and ferroptosis^30^. Ferroptosis is a form of oxidative, non-apoptotic, iron-dependent, cell death that is triggered by excessive lipid peroxide levels **(Figure 4A)**^20,31^. While GOT1 inhibition does not induce ferroptosis, our data suggest it may predispose PDA cells to ferroptosis.

**Figure 4.**
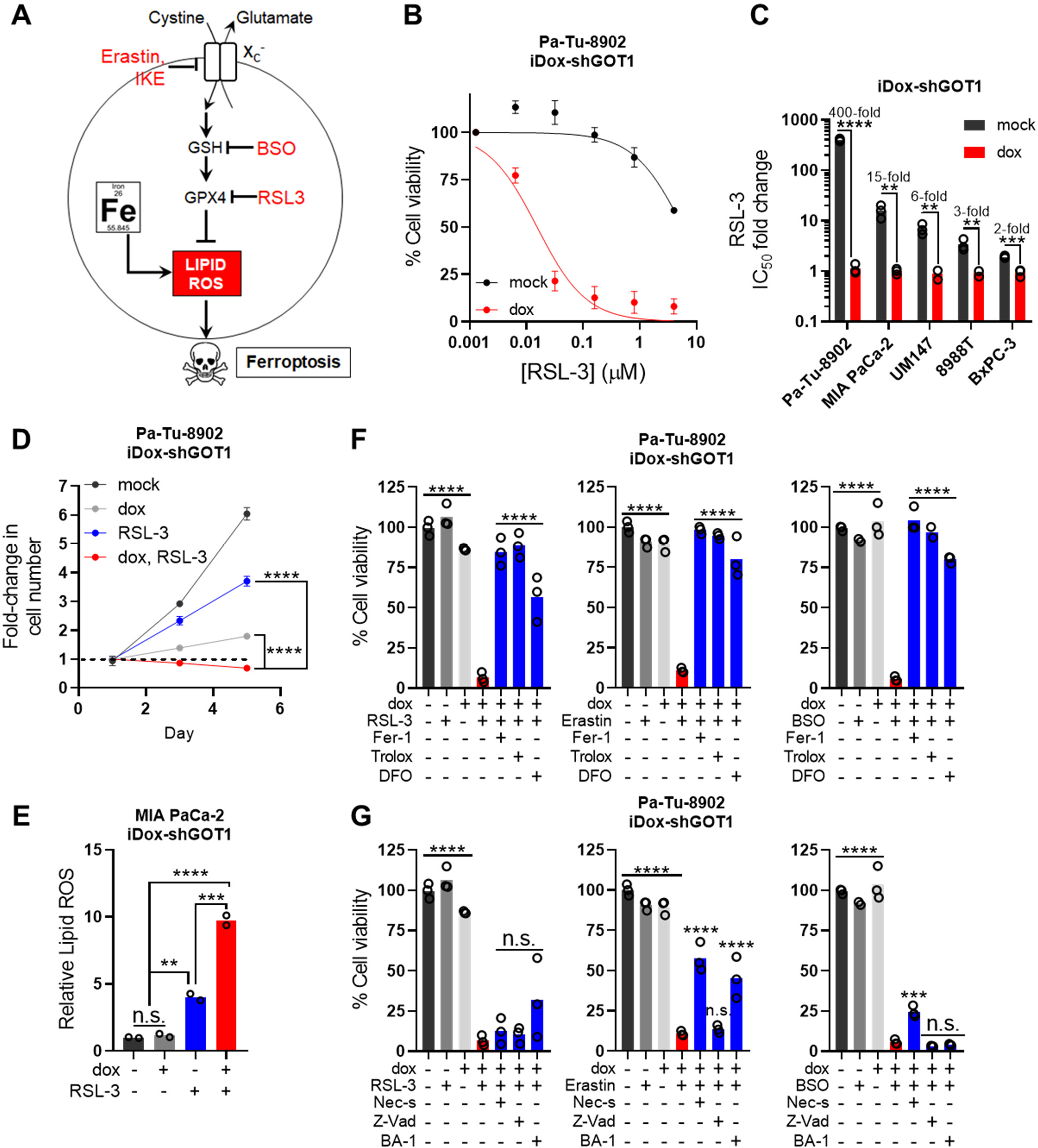
GOT1 suppression sensitizes PDA to ferroptosis. **A**) Schematic of the GPX4 arm of ferroptosis. **B**) Cell viability dose-response curves upon 24 hours of RSL-3 treatment following 5 days of GOT1 knockdown (n=3). Viability is normalized to a 0.1% DMSO vehicle control. **C**) EC_50_ values relative to +dox for 5 iDox-shGOT1 PDA cell lines following 24 hours of RSL-3 treatment (n=3). **D**) Fold change Pa-Tu-8902 iDox-shGOT1 cell numbers following 5 days of GOT1 knockdown and 24 hours treatment with the indicated conditions. 32nM of RSL-3 was administered on day 1. Cell numbers are normalized to day 1 for each condition (n=3). -/+ dox conditions (grey and light grey) contained a 0.1% DMSO vehicle control. **E**) Fold change in viable MIA PaCa-2 iDox-shGOT1 cells positive for C-11 BODIPY, a dye sensitive to lipid ROS, following 5 days of GOT1 knockdown. Cells were treated with the indicated conditions for 6 hours prior to measurements: vehicle (0.1% DMSO) -/+ dox (black and grey), 1μM RSL-3. Data are normalized to the –dox and vehicle-treated condition (n=2). **F**) Cell viability of Pa-Tu-8902 iDox-shGOT1 cultured in vehicle (0.1% DMSO)-/+ dox (black and light grey), drug (32nM RSL-3, 750nM Erastin, 40μM BSO)-/+ doxycycline (grey and red), or drug and dox (blue) in the presence of lipophilic antioxidants 1μM Fer-1 and 100μM Trolox, or an iron chelator 10μM DFO (deferoxamine). Viability was assessed after 24 hours of treatment for RSL-3 and Erastin conditions and 72 hours for BSO treatment conditions. GOT1 was knocked down for 5 days prior to treatment. Data are normalized to the-dox and vehicle treated control (n=3) **G**) Cell viability of Pa-Tu-8902 iDox-shGOT1 cultured in vehicle (0.1% DMSO)-/+ dox (black and light grey), drug (32nM RSL-3, 750nM Erastin, 40μM BSO)-/+ dox (grey and red), or drug and dox (blue) in the presence of 10μM Necrostatin-1 (Nec-s, apoptosis inhibitor), 50 μM ZVAD-FMK (Z-Vad, apoptosis inhibitor), or 1 nM bafilomycin A1 (BA-1, autophagy inhibitor). Viability was assessed after 24 hours of treatment for RSL-3 and Erastin conditions and 72 hours for BSO treatment conditions. GOT1 was knocked down for 5 days prior to treatment. Data are normalized to the-dox and vehicle treated control (n=3). Error bars represent mean ± SD. Two-tailed unpaired T-test or 1-way ANOVA: Non-significant P > 0.05 (n.s. or # as noted), p ≤ 0.05 (*), ≤ 0.01 (**), ≤ 0.001 (***), ≤0.0001 (****). See also **Figure S4**.

To investigate whether GOT1 can sensitize PDA to ferroptosis, we first examined the combinatorial effect of GOT1 knockdown together with RSL-3, a covalent inhibitor of GPX4 and direct inducer of ferroptosis^30^. RSL-3 in combination with GOT1 knockdown was substantially more potent than as a single agent **(Figure 4B)**, and this effect was evident across a panel of PDA lines **(Figures 4C and S4A,B)** and independent of dox effects **(Figure S4C)**. Pa-Tu-8902 is among the most resistant PDA cell lines to single agent GPX4 inhibitors while Mia PaCa-2 was among the most sensitive **(Figures S4A,B)**, in agreement with previous reports suggesting response to GPX4 inhibitors can be predicted by epithelial and mesenchymal markers^32^. Low concentrations of RSL-3 are cytostatic. When combined with GOT1 inhibition, we observed a cytotoxic response **(Figures 4D and S4D)**, suggesting GOT1 inhibition may impose GPX4 dependence in a manner distinct from cell-state and in parallel with GPX4 function.

We then measured the lipid peroxide levels with the C11-BODIPY lipid peroxidation sensor to investigate how inhibition of GPX4 and GOT1 affected lipid peroxidation. While the effect of GOT1 inhibition on lipid ROS induction was modest GOT1 inhibition combined with RSL-3 or erastin substantially upregulated lipid ROS **(Figures 4E and S4E)** and the effect could be reversed through co-treatment with the lipophilic antioxidant ferrostatin-1 (Fer-1) **(Figure S4F)**. Next, we examined whether cell death could be prevented by co-treatment with agents that relieve lipid peroxidation or chelate iron^20,31,33^.

Co-treatments with Fer-1 and Trolox, prevented cell death induced by GOT1 knockdown combined with erastin, BSO or RSL-3 **(Figures 4F and S4H,I)**, while treatment with the iron chelator deferoxamine (DFO), provided substantial protection **(Figure 4F)**. To rule out the possibility that GOT1 was sensitizing PDA to alternative mechanisms of cell death, namely apoptotic, necrotic, or autophagic cell death, we co-treated GOT1 knockdown with well-characterized inhibitors of these cell death pathways. Indeed, the addition of a pan-caspase inhibitor (Z-VAD-FMK), RIPK-1 inhibitor (Necrostatin-1), or lysosomal acidification inhibitor (Bafilomycin A1) offered limited protection from cell death compared with lipophilic antioxidants or iron chelation **(Figures 4G and S4H,I)**, suggesting ferroptosis is the predominant mechanism of cell death. Next, we explored triggering ferroptosis by (–)–FINO_2_ which causes iron oxidation and indirectly inhibits GPX4 activity^34^. Indeed, GOT1 suppression sensitized PDA to (–)–FINO_2_ in an additive manner **(Figure S4J)**. Overall, our data demonstrate GOT1 inhibition primes PDA for ferroptosis.

### GOT1 inhibition primes PDA for ferroptosis by promoting labile iron release in response to metabolic stress

Ferroptosis is driven by oxidation of polyunsaturated fatty acids in the cell membrane, a process that is catalyzed by iron. Thus, ferroptosis is coupled to the cell’s metabolic state^31^. For example, cells can be primed for ferroptosis by enriching polyunsaturated acid composition in cell the membrane in a HIF-2α dependent manner, by depleting essential co-factors for GPX4-^30^ or ferroptosis suppressor protein 1 (FSP1)-mediated^35,36^ lipid antioxidant activity, or by increasing intracellular free iron levels^37^. Because our data indicated that GOT1 inhibition sensitizes PDA to ferroptosis in a manner that is independent of epithelial or mesenchymal cell state, we sought to identify metabolic features that would promote ferroptosis sensitivity in addition to modulating NADPH and GSH levels **(Figure S5A)**.

First, we examined the possibility that GOT1 could be altering phospholipid composition. HIF-2α stabilization has been recently shown to promote a ferroptosis primed cell-state by selectively enriching for polyunsaturated acids in clear-cell carcinomas^38^. This vulnerability was mediated by hypoxia-inducible lipid droplet-associated protein (HILPDA). In our PDA models, both HIF-2α levels and HILPDA expression were unchanged in response to GOT1 knockdown **(Figures S5B-D)**. Next, we examined the possibility that GOT1 inhibition was priming cells to ferroptosis by increasing intracellular iron levels. First, we tested whether GOT1 inhibition would sensitize PDA to perturbations in intracellular iron. We began by testing whether GOT1 knockdown rendered PDA susceptible to increased iron loads using ferric ammonium citrate (FAC) **(Figures 5A and S5E)**. This is consistent with the idea that GOT1 inhibition promotes an oxidized state by depleting both NADPH and GSH while increasing intracellular ROS^8,18^. This oxidized metabolic state primes cells for iron-mediated lipid peroxidation.

**Figure 5.**
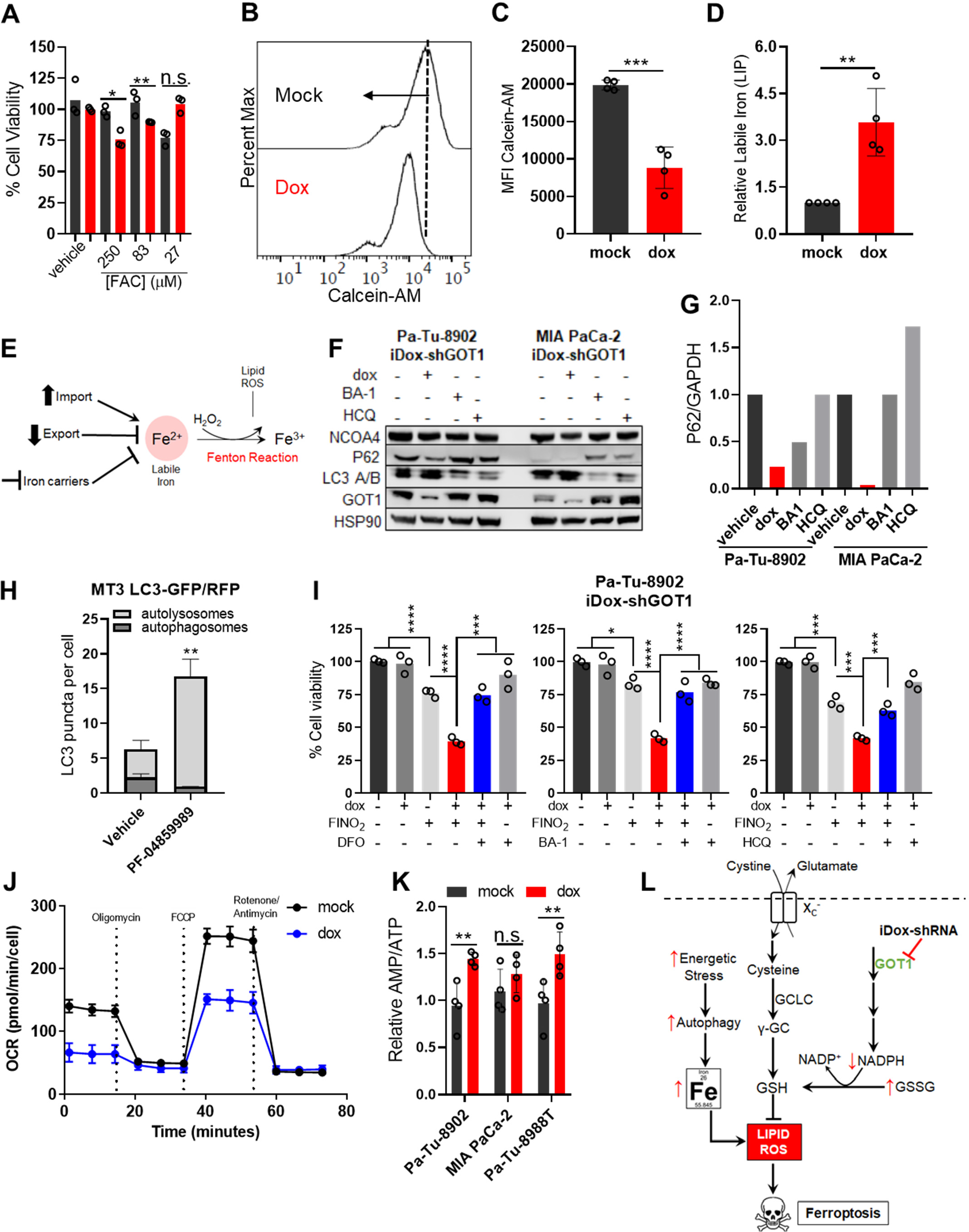
GOT1 inhibition promotes autophagic labile iron release. **A**) Cell viability dose-response curves upon 72 hours of FAC treatment following 5 days of GOT1 knockdown (n=3). Viability is normalized to a 0.1% DMSO vehicle control. **B-D**) Calcein-AM histogram (**B**), mean fluorescence intensity (MFI) (**C**), and relative labile iron (**D**) in Pa-Tu-8902 iDox-shGOT1 cells after 5 days of GOT1 knockdown (n=4). **E**) Mechanisms regulating intracellular iron levels and the Fenton Reaction. **F**) Immunoblot analysis of autophagy markers NCOA4, LC-3 A/B, and P62 in Pa-Tu-8902 and MIA PaCa-2 iDox-shGOT1 cells following 5 days of knockdown. **G**) Quantification of P62 from (F) normalized to the GAPDH loading control. **H**) Quantification of autophagic flux in MT3 cells treated with vehicle or PF-04859989 (n=3) frames. **I**) Cell viability of Pa-Tu-8902 iDox-shGOT1 after 24 hour treatments with 10 μM (–)–FINO_2_ as a single agent, or combined with dox, or dox plus 10 μM deferoxamine (DFO), 8nM bafilomycin A1 (BA-1), or 1.5 μM Hydroxychloroquine (HCQ) (n=3). **J**) Mitostress assay measuring OCR in Mia PaCa-2 iDox-shGOT1 cells (n=3). **K**) AMP/ATP ratio measured by liquid-chromatography tandem mass spectrometry, normalized to - dox (n=4). **L**) GOT1 inhibition promotes the release of labile iron through autophagy in response to metabolic stress and sensitizes PDA to ferroptosis. Error bars represent mean ± SD. Two-tailed unpaired T-test or 1-way ANOVA: Non-significant P > 0.05 (n.s. or # as noted), P ≤ 0.05 (*), ≤ 0.01 (**), ≤ 0.001 (***), ≤0.0001 (****). See also Figure S5.

We then used Calcein-AM dye, a fluorescein-derived probe that is quenched when bound to ferrous iron (Fe^2+^)^39^, to directly assess how GOT1 knockdown was impacting cellular iron levels. Calcein-AM staining of GOT1 proficient cells defined basal fluorescence. Interestingly, GOT1 knockdown cells shifted fluorescence distribution to lower intensity, indicating labile iron pools were increased following GOT1 knockdown **(Figures 5B-D and S5F,G)**. Iron levels can be altered by downregulating iron efflux, upregulating iron uptake, or promoting the degradation of intracellular iron carriers— ferritin or heme **(Figure 5E)**. While previous studies suggest 2017), GOT1 knockdown did not upregulate expression of iron transport proteins (*SLC40A1* and *TFRC*) **(Figures S5H,I)**. Moreover, expression of heme oxygenase 1 (*HMOX1*), which releases labile iron through the degradation of heme was unaltered in Pa-Tu-8902. It was however modestly upregulated in Mia PaCa-2 **(Figure S5I)**, suggesting this mechanism may contribute to increasing intracellular iron pools in some PDA specimens.

Autophagy is a catabolic process that protects cells from metabolic stress induced by nutrient deprivation^40^. Moreover, autophagy is heavily utilized by pancreatic cancers^6,7,41,42^. Indeed, GOT1 knockdown upregulated autophagic flux, indicated by the increase in LC3-B **(Figure 5F)**, consistent with previous work in osteosarcoma cells^43^. Moreover, GOT1 inhibition led to decreased p62, which is selectively degraded during autophagy **(Figure 5F,G)**, and promoted the enrichment of autolysosomes **(Figures 5H and S5J)**. Because GOT1 inhibition did not upregulate the expression of iron importers or exporters to regulate intracellular iron **(Figures S5H,I)**, we wondered if cells may be liberating ferritin-bound iron. transferrin import is required for ferroptosis^44^ and upregulation of transferrin through the iron starvation response can promote ferroptosis^37^.

The majority of stored iron is bound by ferritin and can be released via NCOA4-dependent autophagy, in a process termed ferritinophagy^16^. NCOA4 is autophagosome cargo receptor that binds to the ferritin heavy chain sequestering ferritin for degradation by the autolysosome to release labile iron^16^. Upregulated autophagic flux **(Figure 5F-H)** was matched by NCOA4 expression **(Figure 5F)**, suggesting the changes in labile iron pools may occur through ferritinophagy. Ferritinophagy can indirectly promote ferroptosis by releasing free iron required for lipid peroxidation. Moreover, knockdown of NCOA4 inhibits ferroptosis in HT-1080 fibrosarcoma and in Panc-1, a PDA cell line, potentially by reducing labile iron pools^45,46^. Importantly, supplementing iron chelators (DFO) or inhibitors of lysosomal acidification, [bafilomycin A1 (BA-1) or Hydroxychloroquine (HCQ)] prevented the additive effect of GOT1 inhibition and (–)– FINO_2_ **(Figure 5G)**. Because (–)–FINO_2_ oxidizes iron and since GOT1 inhibition liberates iron, these data support the model that GOT1 promotes labile iron through ferritinophagy.

Autophagy is regulated by cellular energetic and nutrient status. Accordingly, we hypothesized that GOT1 inhibition may induce autophagy in response to metabolic stress. First, using bioenergetic profiling with the Seahorse instrument, we found that GOT1 knockdown lowered basal, maximal, and spare respiratory capacities **(Figure 5J)**, indicating decreased mitochondrial fitness. Further, using LC/MS we found that PDA were experiencing energetic stress, marked by an upregulated AMP/ATP ratio **(Figures 5K and S5K)**. Iron uptake is regulated to support adaption to high oxygen conditions^37^ and to support OxPHOS by supplying metabolically active iron for iron-sulfur cluster and heme biogenesis^47^. Thus, PDA cells may liberate iron as support mitochondrial metabolism following GOT1 loss; however, this results at the cost of rendering PDA susceptible to oxidative assault. Overall, our data suggest that GOT1 primes ferroptosis by promoting the autophagic release of iron in response to metabolic stress **(Figure 5L)**.

## Discussion

In this study we report that the non-canonical malate-aspartate shuttle function is a key contributor to the metabolic fidelity of PDA cells by maintaining NADPH pools and mitochondrial anaplerosis. Inhibition of GOT1 suppresses the growth of numerous PDA cell lines, primary culture models, and xenograft tumors, while rendering cells susceptible to ferroptosis. Ferroptosis could be triggered by inhibition cystine import, glutathione synthesis, or GPX4 in synergy with GOT1 **(Figures 2-4)**, which we ascribe to the promotion of intracellular iron levels through the degradation of ferritin along with the suppression of NADPH.

The dependency on exogenous cystine in GOT1 knockdown cells was identified using a synthetic lethal chemical screening strategy **(Figures 2A-C and S2A-B)**. We then demonstrated that exogenous cystine through system xC^-^ enables *de novo* GSH biosynthesis, which is required to compensate for decreased GSH availability. When GOT1 is inhibited, NADPH availability is decreased and GSH cannot be sufficiently regenerated by reducing GSSG. This metabolic rewiring could be exploited by dietary means, which has a major influence on the nutrient composition within pancreatic tumors^24^. This result adds to a growing body of literature indicating how the metabolic environment can influence sensitivity to therapy^48,49^.

The inhibition of *de novo* GSH biosynthesis with BSO also potentiated tumor inhibition by GOT1. BSO has been used in clinical trials and is tolerated in patients^28,29^. Previous efforts with BSO have indicated that many tumor types employ compensatory mechanisms to tolerate glutathione inhibition^27,50^. Indeed, we too have observed that inhibition of *de novo* glutathione biosynthesis is not sufficient to induce ferroptosis in PDA. This points to GOT1 inhibition as a logical combination therapy to enhance therapeutic efficacy of BSO. To this end, we and others engaged in drug discovery campaigns to develop small inhibitors of GOT1^10,11,12,13^. Among these, we found that the KATII inhibitor PF-04859989 also has potent GOT1 inhibitory activity^13^, and we apply that herein as a tool compound. Future studies are yet required to improve the drug like properties of this molecule for in vivo studies.

Reduced glutathione is a co-factor for GPX4 that is used to maintain lipid antioxidant function. It is also conceivable that the non-canonical malate-aspartate shuttle may supply NADPH to FSP1, which reduces the co-factor CoQ10 to inhibit lipid peroxides in parallel with GPX4^35,36^. This could also account for how GOT1 inhibition sensitizes PDA to ferroptosis **(Figures 3-5)**. These results would be consistent with previous studies demonstrating GOT1 inhibition can induce ROS^8,12^ and radiosensitize PDA both *in vitro* and *in vivo*^18^. Future studies will be required to examine the role of FSP1 and potential interactions with the GOT1 pathway in PDA.

In line with the name, ferrous iron is required for ferroptosis by contributing to the oxidation of membrane PUFAs, either as free iron or as a co-factor for lipoxygenase enzymes. Iron levels can be altered by upregulating iron import, downregulating iron export, or degrading iron carriers. Our studies indicate that GOT1 knockdown promotes the release of intracellular iron, and sensitized PDA to the deleterious effects of iron oxidation and iron loading. GOT1 knockdown did not alter the expression of iron uptake or efflux proteins. Rather, GOT1 knockdown leads to a catabolic state marked by diminished mitochondrial activity and elevated AMP/ATP. To support metabolic homeostasis, GOT1 knockdown cells activate autophagy, a process that enables cells to degrade cellular components through the lysosome and recycle these to adapt to nutrient stress. Concurrently, autophagy in these NCOA4 expressing PDA cells also leads to the turnover of ferritin-bound iron via ferritinophagy^16^. This leads to higher levels of free iron, and by extension, promotes susceptibility to ferroptosis. It is not entirely clear why PDA cells would release free iron through ferritinophagy upon GOT1 knockdown. It has been proposed by that labile iron, which is metabolically active and can be utilized by iron-containing enzymes, is released to support DNA synthesis, epigenetic modification, and the biogenesis of iron-sulfur clusters to support mitochondrial metabolism^47^. Our data are consistent with a model whereby cells can employ ferritinophagy to release iron for metabolic demands, but with the cost of rendering cells susceptible to ferroptosis.

The role of GOT1 in ferroptosis has been the subject of previous study in several other tumor types. Our data lie in contrast to some previous work, which have suggested that GOT1 inhibition by genetic or pharmacological means in other tumor types protects cells from ferroptosis by blocking mitochondrial metabolism^20,45,51^. Recent studies suggest the genotype^32^, nutrient environment^52^, tissue of origin^30,38^, and cell-autonomous metabolism^35-37^ are major drivers of ferroptosis sensitivity. The differences emerging from these studies likely reflect the incomplete understanding of how these various factors dictate sensitivity to ferroptosis. Thus, our study provides clarity regarding the metabolic regulation of ferroptosis in PDA.

### Significance

PDA is a notoriously deadly disease with a ten percent 5-year survival owing in largely to a lack of effective therapeutic options. PDA cells rewire metabolism to survive and proliferate with a nutrient-deprived and metabolically harsh environment. This study not only demonstrates that the GOT1 non-canonical malate-aspartate pathway is essential for maintaining biosynthetic and antioxidant fidelity in pancreatic cancer, it also illustrates that promoting a catabolic, oxidative cell state leads to the autophagy-dependent release of labile iron. This sensitizes pancreatic cancer to ferroptosis by pharmacological or dietary means. These applications represent alternative contexts for the repurposing clinical agents and could reveal novel therapeutic strategies that selectively exploit the unique metabolic demands of pancreatic tumors.

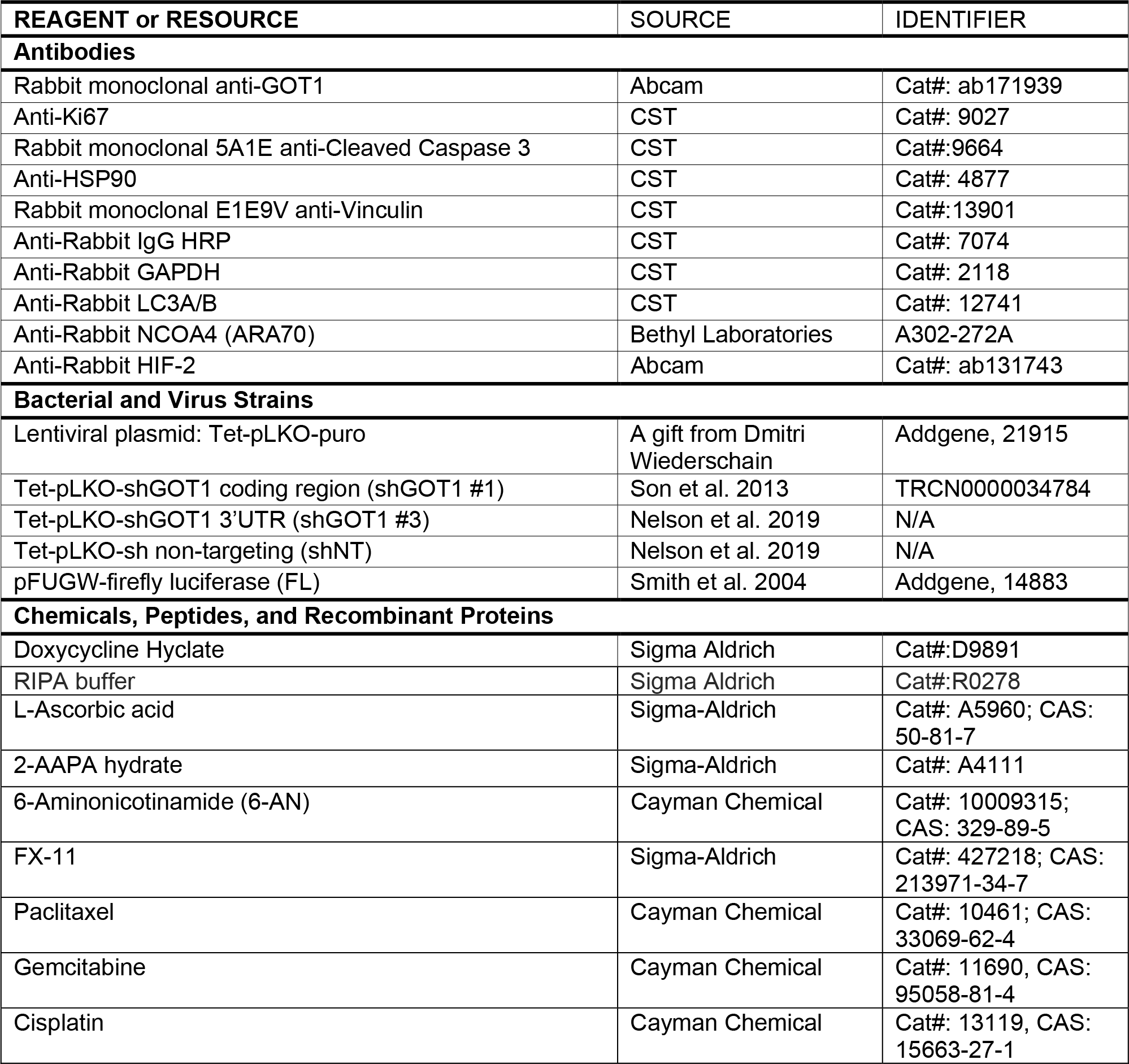
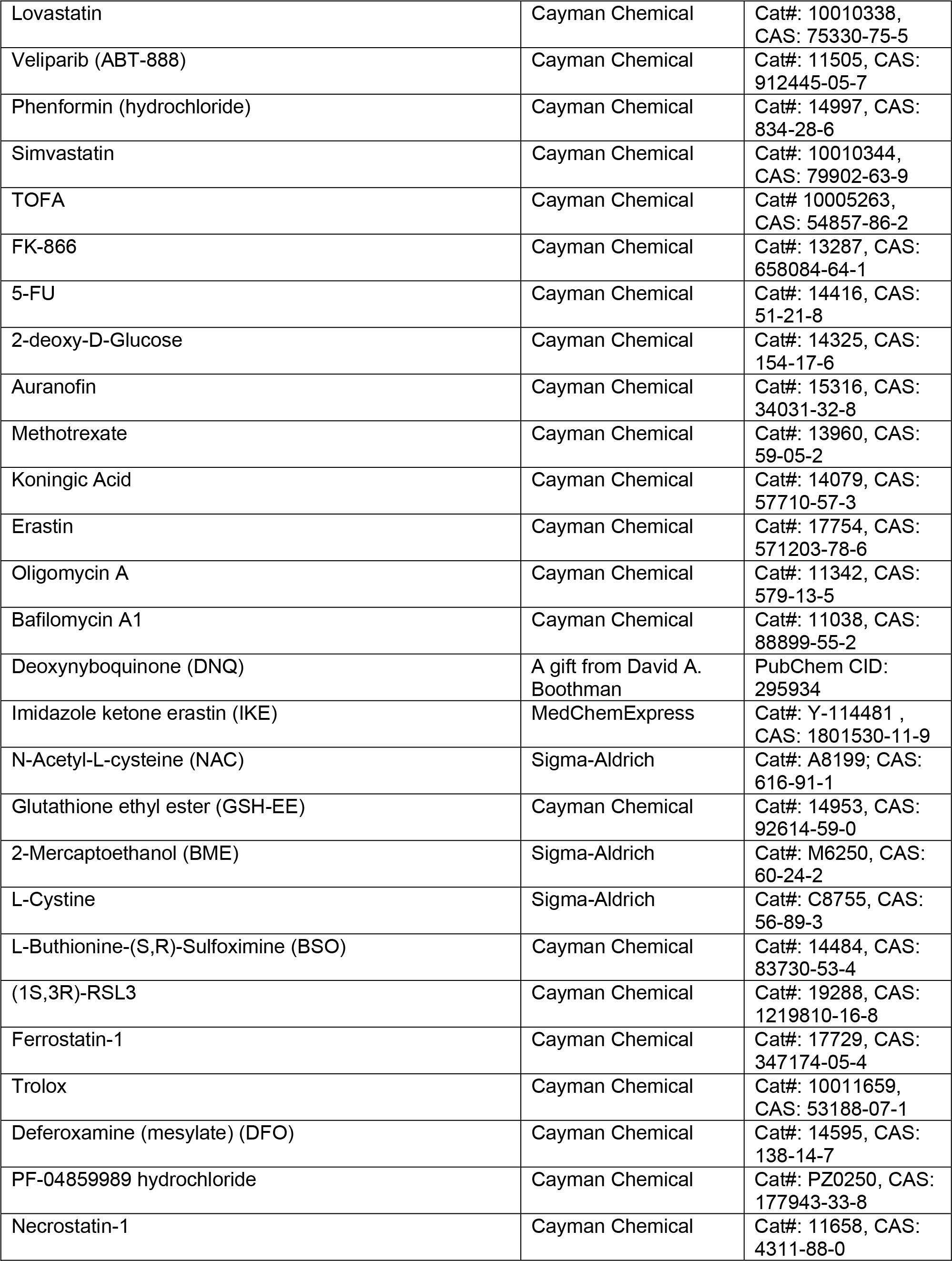
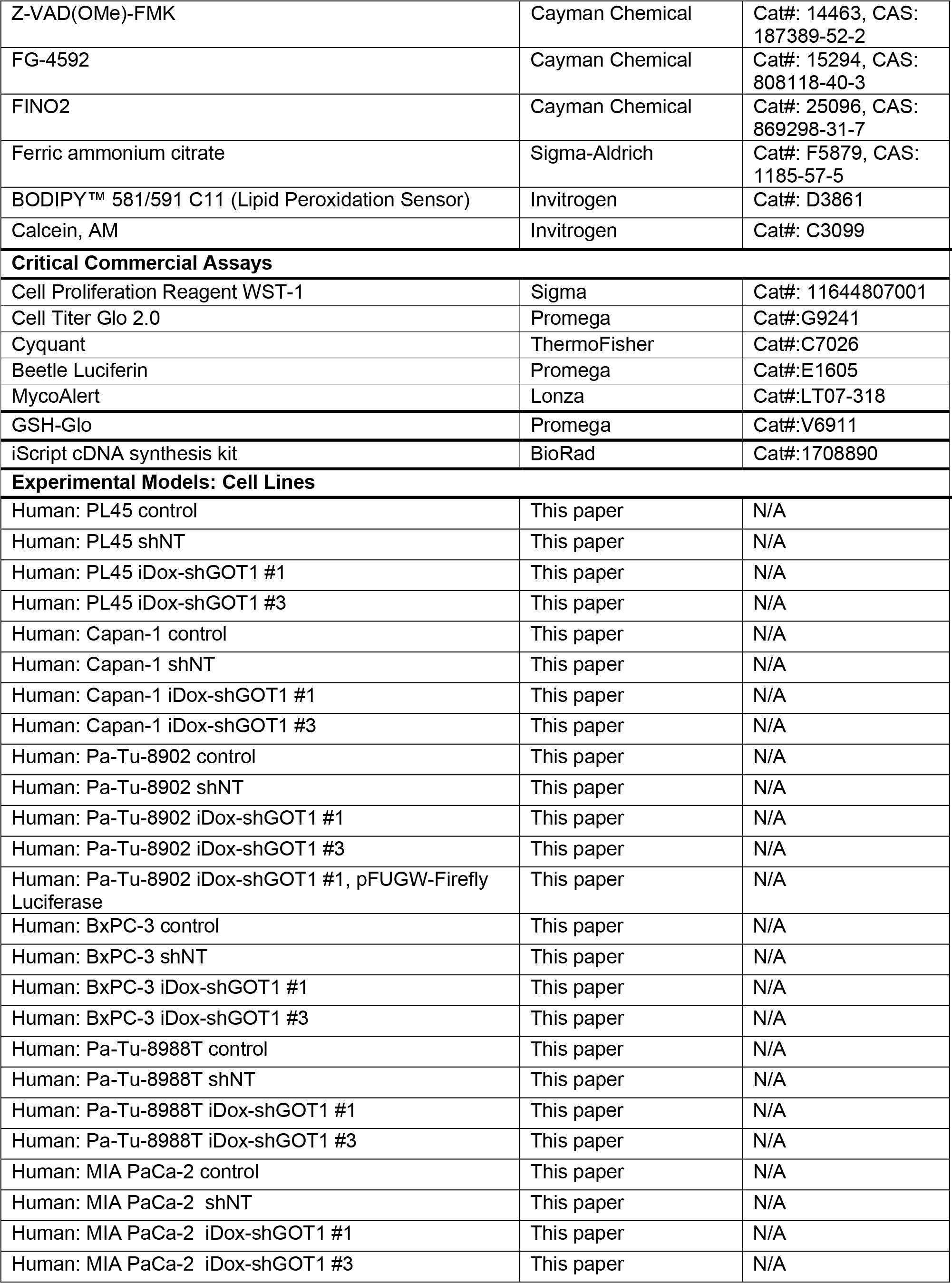
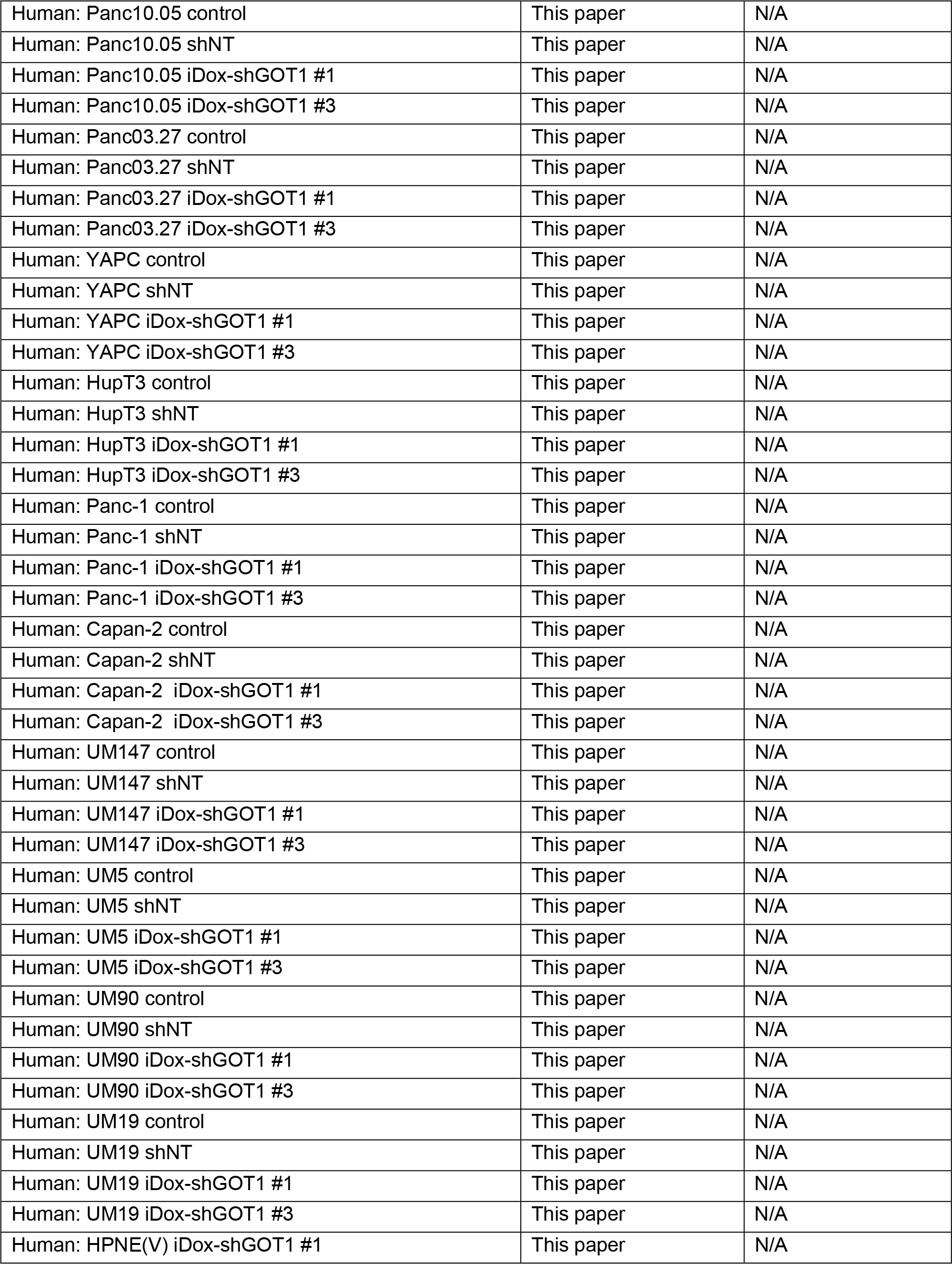
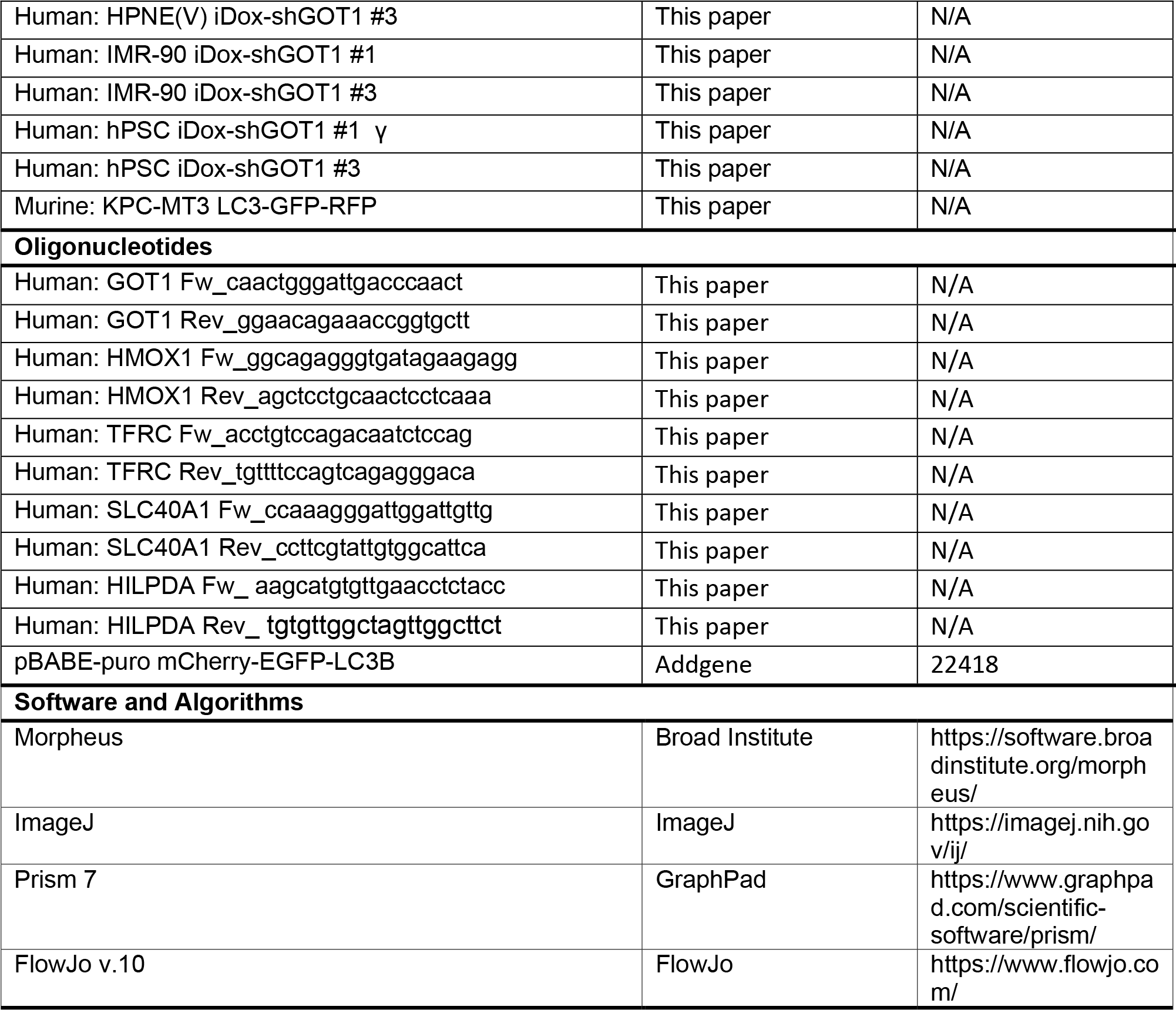
KEY RESOURCES TABLE

### Contact for Reagent and Resource Sharing

Further information and requests for resources and reagents should be directed to and will be fulfilled by the Lead Contact, Costas A. Lyssiotis.

### Cell Culture

PL45, Capan-1, BxPC-3, MIA PaCa-2, Panc10.05, Panc03.27, PANC-1, Capan-2, HPNE (V), IMR-90 were obtained from ATCC. Pa-Tu-8902, Pa-Tu-8988T, YAPC, and Hup T3 were obtained from DSMZ. Human pancreatic stellate cells (hPSC) were a generous gift form Rosa Hwang (Hwang et al., 2008). The UM PDA primary cell cultures (UM147, UM5, UM90, and UM19) were obtained from surgically-resected samples and established through murine xenograft (Li et al., 2007). KPC-MT3 murine PDA cell lines were a generous gift from Dr. David Tuveson. All commercial cell lines and UM PDA primary cultures were validated by STR profiling and tested negative for mycoplasma infection (Lonza, LT07-701). Cells were maintained under standard conditions at 37°C and 5% CO2. Cells were grown either in regular DMEM (GIBCO, #11965) or RPMI (GIBCO, #11875), or in DMEM without cystine (GIBCO, #21013024) or RPMI (GIBCO, A1049101) supplemented with 10% FBS (Corning, 35-010-CV) unless otherwise indicated. Cultures involving inducible short-hairpin mediated knockdown were supplemented with doxycycline-hyclate (Dox) at 1μM/mL (Sigma, D9891) for 5 days prior to experiments.

KPC-MT3 LC3 tandem GFP-RFP cells were established through transfection with pBABE-puro mCherry-EGFP-LC3B (Addgene plasmid #22418) followed by selection and single cell cloning. For autophagic flux quantification, cells were seeded into 4 well chamber slides (Corning, 354104) and fixed in 4% paraformaldehyde (ThermoFisher, 28908) 24 hours following seeding. Coverslips were mounted in DAPI containing mounting solution (Life Technologies P36931). Cells were imaged using a Nikon A1 Confocal in FITC, RFP and DAPI channels. The ratio of red:yellow puncta was determined by counting puncta using the particle analysis in imageJ.

### Lentiviral-mediated shRNA Transduction

Parental PDA cell lines were transduced with lentivirus containing short hairpin RNA plasmids at optimized viral titers. Stable cell lines were established post-puromycin selection.

### Clonogenic assays

Cells were plated in a 6-well plate in biological triplicates at 300-600 cells per well in 2mL of media. Dox-media were changed every 2 days. Assays were concluded after 10-15 days by fixing in −20°C cold 100% methanol 10 min and staining with 0.5% crystal violet 20% methanol solution for 15 min. Colonies were quantified using ImageJ or manually counted.

### Cell proliferation assays

Cells were seeded in a 96-well plate at 1,000 cells per well in 0.1mL of media. Indicated treatments were applied the subsequent day. Media was changed every 2 days. At the indicated time points, media was aspirated and frozen. 100 μL of CyQUANT (Invitrogen, C7026) to each well for measurements. 10μL of WST-1 reagent directly to the culture media (Sigma, #11644807001). Relative proliferation was determined by the fluorescence intensity at 530nm for CyQuant or 450nm for WST-1 using a SpectraMax M3 plate reader.

### Cell viability assays

Cells were plated in a 96-or 384-well plate format at 1,000 cells per well Cells were allowed to seed overnight, then treated with compounds at indicated concentrations and for indicated lengths of time. All viability assays utilized the Cell-Titer-Glo 2.0 reagent (Promega, G9243) according to the manufacturer’s instructions. Media was aspirated followed by the addition of 100 μL of Cell-Titer-Glo 2.0 reagent to each experimental well. Plates were gently agitated for 10 minutes to promote adequate mixing. Luminescence was subsequently measured using a SpectraMax M3 plate reader.

### Quantitative RT-PCR

Total RNA was extracted using the RNeasy Mini Kit (Qiagen, 74104) and reverse transcription was performed from 2 μg of total RNA using the iScript cDNA synthesis kit (BioRad,1708890) according to the manufacturer’s instructions. Quantitative RT–PCR was performed with Power SYBR Green dye (Thermo, 4367659) using a QuantStudio 3 System (Thermo). PCR reactions were performed in triplicate and the relative amount of cDNA was calculated by the comparative *C_T_* method using an *RPS21* as an endogenous control. RT-PCR was performed in a least 3 biological replicates.

### Detection of Reactive Oxygen and Labile Iron by Flow cytometry

Cells were plated in 6-well plates two days before incubation with indicated treatments. Cells were then washed twice with 1x PBS, and stained for 20-30 (Invitrogen, C1430) minutes with 2μM C11-BODIPY (Invitrogen, D3861) or for 10 minutes with 0.2μM Calcein-AM (Invitrogen, C1430) in phenol red-free DMEM. Cells were co-stained with Sytox-blue (Invitrogen, S34857) to account for cell viability. Following staining, cells were washed twice with PBS, trypsinized (0.25%, Life Technologies, 25200-056), and naturalized with pure FBS at a 1:1 volume. Cells were then collected in 500uL PBS, and moved to round bottom 96-well plates, on ice, for measurements. A minimum of 8000 cells were analyzed per condition. C11-BODIPY and Calcein-AM signals were analyzed in the FITC channel, while Sytox-blue was analyzed in the DAPI channel on a ZE5 Cell analyzer (Bio-Rad). Analysis of data was performed using FlowJo v.10 software. See **Supplementary Figures 6 and 7** for representative gating. Relative labile iron levels were calculated based on the ratio of Calcein-AM mean fluorescence intensity (MFI) of control vs. dox-treated samples.

### Xenograft Studies

Animal experiments were conducted in accordance with the Office of Laboratory Animal Welfare and approved by the Institutional Animal Care and Use Committees of the University of Michigan. NOD scid gamma (NSG) mice (Jackson Laboratory, 005557), 6-8 or 8-10 weeks old of both sexes, were maintained in the facilities of the Unit for Laboratory Animal Medicine (ULAM) under specific pathogen-free conditions. Stable PDA cell lines containing a dox-inducible shRNA against GOT1 were trypsinzied and suspended at 1:1 ratio of DMEM (Gibco, 11965-092) cell suspension to Matrigel (Corning, 354234). 150-200 μL were used per injection. For subcutaneous xenograft studies, 0.5×10^6^ cells were implanted into the lower flanks. Doxycycline (dox) chow (BioServ, F3949) was fed to the +dox groups. Orthotopic tumors were established by injecting 5×10^4^ Pa-Tu-8902 iDox-shGOT1 #1 pFUGW-Firefly Luciferase into 8-10 week old NSG mice. Cysteine-free chow (LabDiet) was customized from Baker Amino Acid (LabDiet, 5CC7) to remove cysteine and balance protein levels with increased valine and aspartic acid. BSO was delivered in the drinking water at 20 mM. All treatments began on day 7 after implantation.

Subcutaneous tumor size was measured with digital calipers at the indicated endpoints. Tumor volume (V) was calculated as V = 1/2(length x width^2^). Bioluminescence (BLI) of orthotopic tumors were measured via IVIS SpectrumCT (PerkinElmer) following an intraperitoneal injection of 100 μL beetle luciferin (40 mg/mL in PBS stock) (Promega, E1605). BLI was analyzed with Living Image software (PerkinElmer). At endpoint, final tumor volume and mass were measured prior to processing. Tissue was either fixed in zinc formalin fixative (Z-fix, Anatech LTD, #174) for >24 hours for histological and/or histochemical analysis, or snap-frozen in liquid nitrogen then stored at −80°C until metabolite or protein analysis.

### Western blot analysis

Stable shNT and shGOT1 cells were cultured with or without dox media and protein lysates were collected after five days using RIPA buffer (Sigma, R0278) containing protease inhibitor cocktail (Sigma/Roche, 04 693 132 001). Samples were quantified with Pierce BCA Protein Assay Kit (ThermoFisher, 23225). 10 to 40 μg of protein per sample were resolved on NuPAGE Bis-Tris Gels (Invitrogen, NP0336) and transferred to a Immobilon-FL PVDF membrane (Millipore, IPVH00010). Membranes were blocked in 5% non-fat dry milk in distilled H2O prior to incubation with the primary antibody. The membranes were washed with TBS-Tween followed by a 1h exposure to the appropriate horseradish peroxidase-conjugated secondary antibody. The membranes were washed in de-ionized water for 15-30 minutes then visualized using a Bio-Rad ChemiDox MP Imaging System (Bio-Rad, 17001402). The following antibodies were used: anti-aspartate aminotransferase (anti-GOT1) at a 1:1,000 dilution (Abcam, ab171939), 1:1,000 dilution Anti-Rabbit LC3 A/B (CST, 12741), 1:1,000 dilution Anti-Rabbit NCOA4 (Bethyl Laboratories, A302-272A), 1:1,000 dilution Anti-Rabbit Hif-2 (Abcam, ab131743), and loading control vinculin at a 1:1,000 dilution (Cell Signaling, 13901), HSP-90 (Cell Signaling, 4877S), Anti-Rabbit β-Actin (Cell Signaling, 4970L) or GAPDH (Cell Signaling, 2118). Anti-rabbit IgG, HRP-linked (Cell Signaling Technology, 7074) secondary antibody was used at a 1:10,000 dilution.

### Histology

Mice were sacrificed by CO2 asphyxiation followed by tissue harvesting and fixation overnight at room temperature with Z-fix solution (Z-fix, Anatech LTD, #174). Tissues were the processed by using a Leica ASP300S Tissue Processor, paraffin embedded, and cut into 5-μm sections. Immunohistochemistry was performed on Discovery Ultra XT autostainer (Ventana Medical Systems Inc.) and counterstained with hematoxylin. IHC slides were scanned on a Panoramic SCANslide scanner (Perkin Elmer), and then annotation regions encompassing greater than 1mm of tissue were processed using Halo software (Indica Labs).The following antibodies were used for IHC: GOT1 (AbCam, ab171939), Ki-67 (Cell Signaling, 9027), Cleaved Caspase-3 (Cell Signaling, 9664).

### Metabolomics

Targeted metabolomics: Cells were plated at 500,000 cells per well in 6-well plates or ~1.5 million cells per 10 cm dish. At the endpoint, cells were lysed with dry-ice cold 80% methanol and extracts were then centrifuged at 10,000 g for 10 min at 4°C and the supernatant was stored at −80°C until further analyses. Protein concentration was determined by processing a parallel well/dish for each sample and used to normalize metabolite fractions across samples. Based on protein concentrations, aliquots of the supernatants were transferred to a fresh micro centrifuge tube and lyophilized using a SpeedVac concentrator. Dried metabolite pellets were re-suspended in 45 μL 50:50 methanol:water mixture for LC–MS analysis. Data was collected using previously published parameters^53,54^.

The QqQ data were pre-processed with Agilent MassHunter Workstation Quantitative Analysis Software (B0700). Additional analyses were post-processed for further quality control in the programming language R. Each sample was normalized by the total intensity of all metabolites to scale for loading. Finally, each metabolite abundance level in each sample was divided by the median of all abundance levels across all samples for proper comparisons, statistical analyses, and visualizations among metabolites. The statistical significance test was done by a two-tailed t-test with a significance threshold level of 0.05.

### Seahorse Mito Stress Test

MiaPaCa-2 cells were seeded at 2×10^4^ cells/well in 80 μl/well of normal growth media (DMEM with 25 mM Glucose and 2 mM Glutamine) in an Agilent XF96 V3 PS Cell Culture Microplate (#101085-004). To achieve an even distribution of cells within wells, plates were incubated on the bench top at room temperature for 1 hour before incubating at 37°C, 5% CO2 overnight. To hydrate the XF96 FluxPak (#102416-100), 200 μL/well of sterile water was added and the entire cartridge was incubated at 37’C, no CO2 overnight. The following day, one hour prior to running the assay, 60 μL/well of growth media was removed from the cell culture plate and cells were washed twice with 200 μL/well of assay medium (XF DMEM Base Medium, pH 7.4 (#103575-100) containing 25 mM Glucose (#103577-100) and 2 mM Glutamine (#103579-100)). After washing, 160 μL/well of assay medium was added to the cell culture plate for a final volume of 180 μL/well. Cells were then incubated at 37’C, no CO2 until analysis. Also one hour prior to the assay, water from the FluxPak hydration was exchanged for 200 μL/well of XF Calibrant (#100840-000) and the cartridge was returned to 37’C, no CO2 until analysis.

Oligomycin (100 μM), FCCP (100 μM), and Rotenone/Antimycin (50 μM) from the XF Cell Mito Stress Test Kit (#103015-100) were re-constituted in assay medium to make the indicated stock concentrations. 20 μL of Oligomycin was loaded into Port A for each well of the FluxPak, 22 μL of FCCP into Port B, and 25 μL of Rotenone/Antimycin into Port C. Port D was left empty. The final FCCP concentration was optimized to achieve maximal respiration in each condition. The Mito Stress Test was conducted on an XF96 Extracellular Flux Analyzer and OCR was analyzed using Wave 2.6 software. Following the assay, OCR was normalized to cell number with the CyQUANT NF Cell Proliferation Assay (C35006) from Thermo Fisher according to manufacturer’s instructions.

### Statistical analysis

Statistics were performed using GraphPad Prism 7 (Graph Pad Software Inc). Groups of 2 were analyzed using the unpaired two-tailed Student’s t test and comparisons across more than 2 groups were conducted using one-way ANOVA Tukey post-hoc test. All error bars represent mean with standard deviation, unless noted otherwise. A P value of less than 0.05 was considered statistically significant. All group numbers and explanation of significant values are presented within the figure legends.

## Acknowledgements

The authors would like to thank members of the Olive, Shah, and Lyssiotis lab for critical discussion and scientific feedback. D.M.K was supported by a Department of Education GAANN fellowship through The University of Michigan Program in Chemical Biology. B.S.N was supported by Institutional National Research Service Awards through NIDDK (T32-DK094775) and NCI (T32-CA009676). M.P.M. was supported by NCI R01-CA-198074, R01-CA-151588, and an ACS Research Scholar Grant. Y.M.S was supported by NIH grants (R01CA148828 and R01DK095201). C.A.L. was supported by the Pancreatic Cancer Action Network/AACR (13-70-25-LYSS), Damon Runyon Cancer Research Foundation (DFS-09-14), V Foundation for Cancer Research (V2016-009), Sidney Kimmel Foundation for Cancer Research (SKF-16-005), the AACR (17-20-01-LYSS), and an ACS Research Scholar Grant (RSG-18-186-01). M.P.M., Y.M.S., H.C.C., and C.A.L. were supported by the Cancer Center Support Grant (P30 CA046592). M.P.M., H.C.C., and C.A.L. were supported by U01-CA-224145. K.P.O. and C.A.L. were supported by R01CA215607. Metabolomics studies at U-M were supported by DK09715, the Charles Woodson Research Fund, and the UM Pediatric Brain Tumor Initiative.

## Author Contributions

Conceptualization, D.M.K. and C.A.L.; Investigation, D.M.K, B.S.N., L.L., E.L.Y, A.M., G.T., C.J.H., S.A.K., N.C., S.W., J.R., S.S., C.P., and T.G.; Resources, E.C., L.Z., M.A.B., P.S., H.C.C., A.C.A., Z.C.N., J.M.A., M.P.dM., Y.M.S, and K.P.O.; Data Curation, D.M.K., P.S., A.C.A, and L.Z.; Writing – Original Draft, D.M.K. and C.A.L.; Writing – Review & Editing, D.M.K., B.S.N, E.L.Y., Z.C.N., Y.M.S., K.P.O., and C.A.L.; Formal analysis and Visualization, D.M.K; Supervision and Funding Acquisition, C.A.L.

## Supplemental Figure Legends

**Supplementary Figure 1, related to Figure 1.**
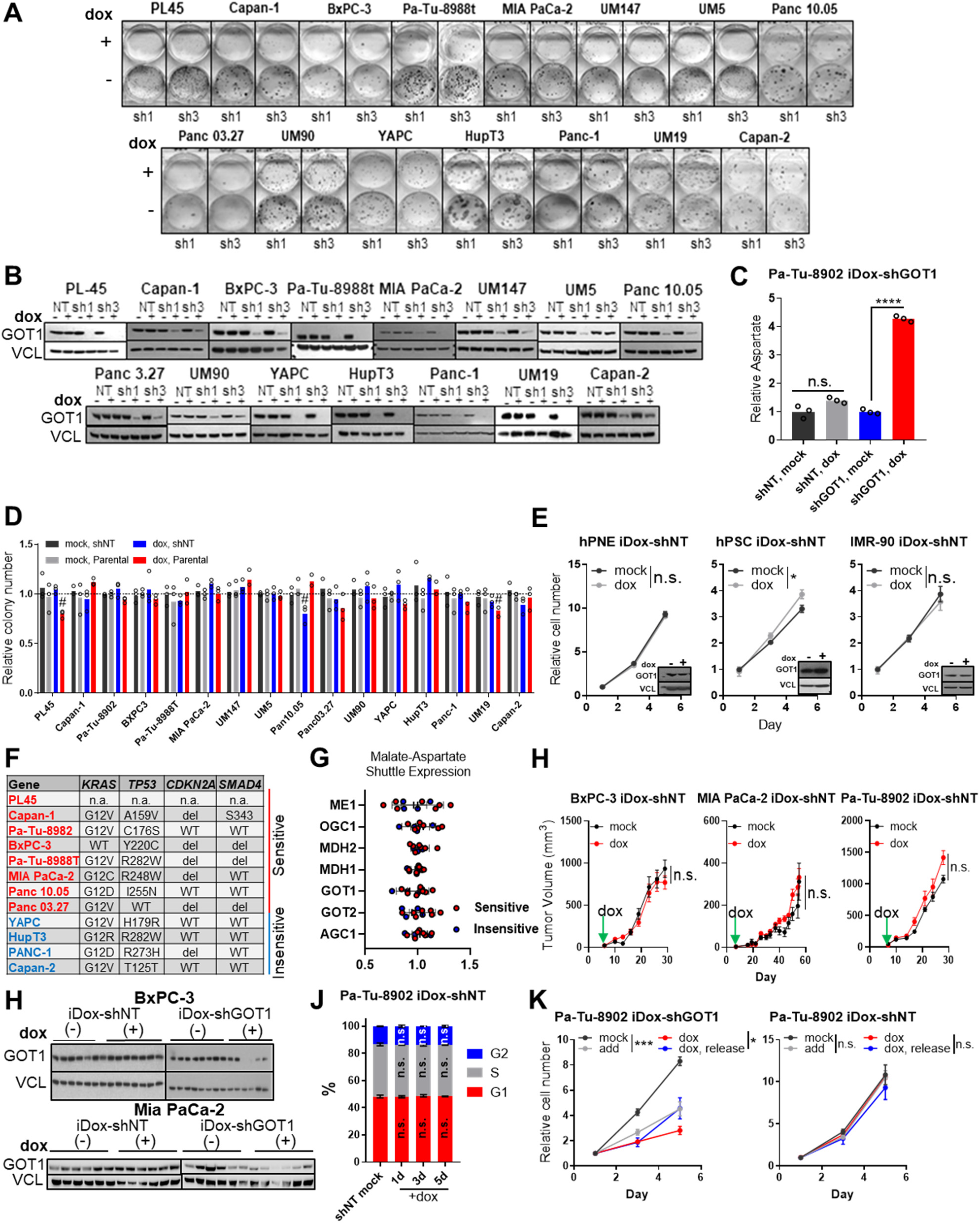
GOT1 dependence varies among pancreatic cancers. **A**) Representative colony formation after 10-15 days (n=3) corresponding to **Figure 1D**. **B**) Immunoblots corresponding to Figure 1D and SFigure 1A. Non-targeting, sh1, and sh3 stable cell lines were treated with dox for 5 days. VCL was used as a loading control. **C**) Intracellular aspartate abundance in Pa-Tu-8902 shNT and sh1 stable cell lines after 5 days of dox-treatment accessed by liquid-chromatography tandem mass spectrometry, normalized to mock (n=3). **D**) Relative colony counts for NT (black/blue) stable cell lines and parental cell lines (grey/red) corresponding to **Figure 1D** after 10-15 days (n=3). **E**) Relative cell number of immortalized non-transformed human cell lines with shNT vectors corresponding to Figure 1E. 1, 3, or 5-day time points were used, normalized to day 1 (n=3). **F**) Common mutations associated with PDA from the Cancer Cell Line Encyclopedia. **G**) Expression (mRNA) of malate-aspartate shuttle enzymes in GOT1 sensitive (n=7) and insensitive (n=4) PDA cell lines from the Cancer Cell Line Encyclopedia. Sensitivity based on data in **Figure 1D**. **H**) Growth of subcutaneous xenograft tumors containing non-targeting (NT) vectors treated with dox (red) or vehicle (black) (n= 6), corresponding to Figure 1F. **I**) Immunoblots for GOT1 corresponding to Figures 1J and SFigure 1H. VCL was used as a loading control. **J**) Cell cycle distribution of Pa-Tu-8902 cells expressing shNT constructs corresponding to **Figure 1K**. **K**) Relative cell number in iDox-shGOT1 or NT cells corresponding to Figure 1H. 1, 3, and 5-day time points were used. Cell counts were normalized to day 1 (n=3). Error bars represent mean ± SD. Non-significant P > 0.05 (n.s. or # as noted), P ≤ 0.05 (*), ≤ 0.01 (**), ≤ 0.001 (***), ≤ 0.0001 (****). See also **Figure 1**.

**Supplementary Figure 2, related to Figure 2.**
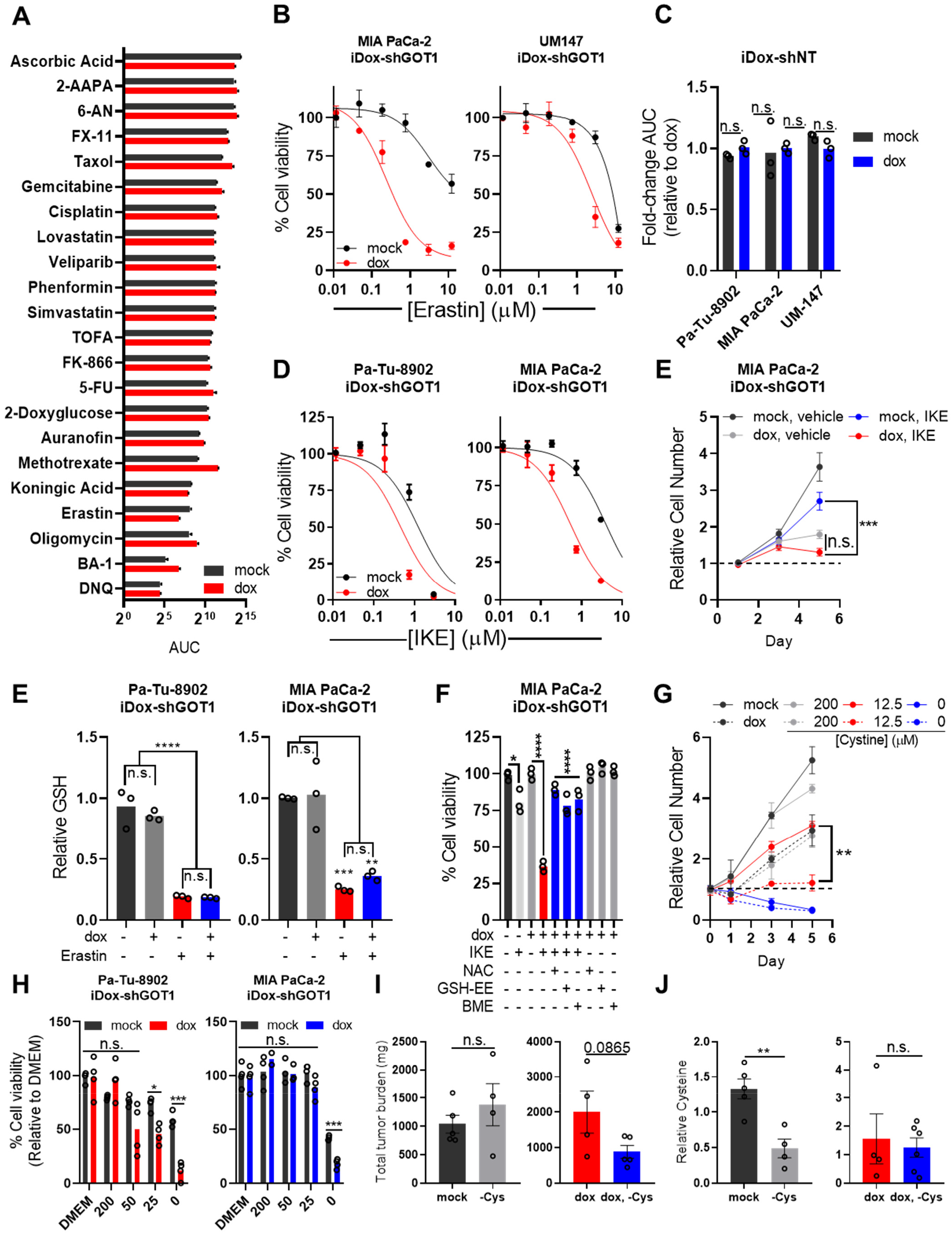
GOT1 knockdown sensitizes PDA to xCT inhibition and cystine deprivation. **A**) Area under the curve (AUC) in cell viability for each compound in **Figure 2B** for mock (black) or knockdown conditions (red) after 72 hours (n=3). **B-D**) Cell viability dose response curves for erastin after 24 hours in iDox-shGOT1 expressing cell lines (**B**) depicted in **Figure 2D**. shNT expressing cell lines were used in (**C**) and the AUC fold-change is presented, while imidazole ketone erastin (IKE) was used in (**D**) (n=3). **E**) Relative MIA PaCa-2 iDox-shGOT1 cell numbers after 5 days of dox treatment with the indicated conditions. 750nM of IKE was administered on day 1 and each condition is normalized to day 1 (n=3). **F**) Relative reduced glutathione levels (GSH) after 5 days of knockdown followed by 750nM of erastin for 6 hours. GSH levels were first normalized to cell viability and then normalized again to vehicle treatment (black and grey) (n=3). **G**) Cell viability of Mia PaCa-2 iDox-shGOT1 after 5 days of dox culture then 750nM IKE co-cultured with the indicated conditions (n=3). **H**) Relative Mia PaCa-2 iDox-shGOT1 cell numbers following 5 days of GOT1 knockdown and the indicated media conditions (n=3). **I**) Cell viability of Pa-Tu-8902 iDox-shGOT1 (red) and Mia PaCa-2 iDox-shGOT1 (blue) following 5 days of dox pre-treatment and 24 hours of the indicated cystine concentrations (n=4). **J**) Total post-treatment tumor burden (mock, n=5), (-Cys, 4), (dox, n=4), and (dox, -Cys, n=5). **K**) Cysteine levels in tumors at endpoint (mock, n=5), (-Cys, 4), (dox, n=4), and (dox, - Cys, n=6). Error bars represent mean ± SD (Figures S2a-h) or mean ± S.E.M (Figures S2i-j). Two-tailed unpaired T-test or 1-way ANOVA: Non-significant P > 0.05 (n.s. or # as noted), P ≤ 0.05 (*), ≤ 0.01 (**), ≤ 0.001 (***), ≤0.0001 (****). See also **Figure 2**.

**Supplementary Figure 3, related to Figure 3.**
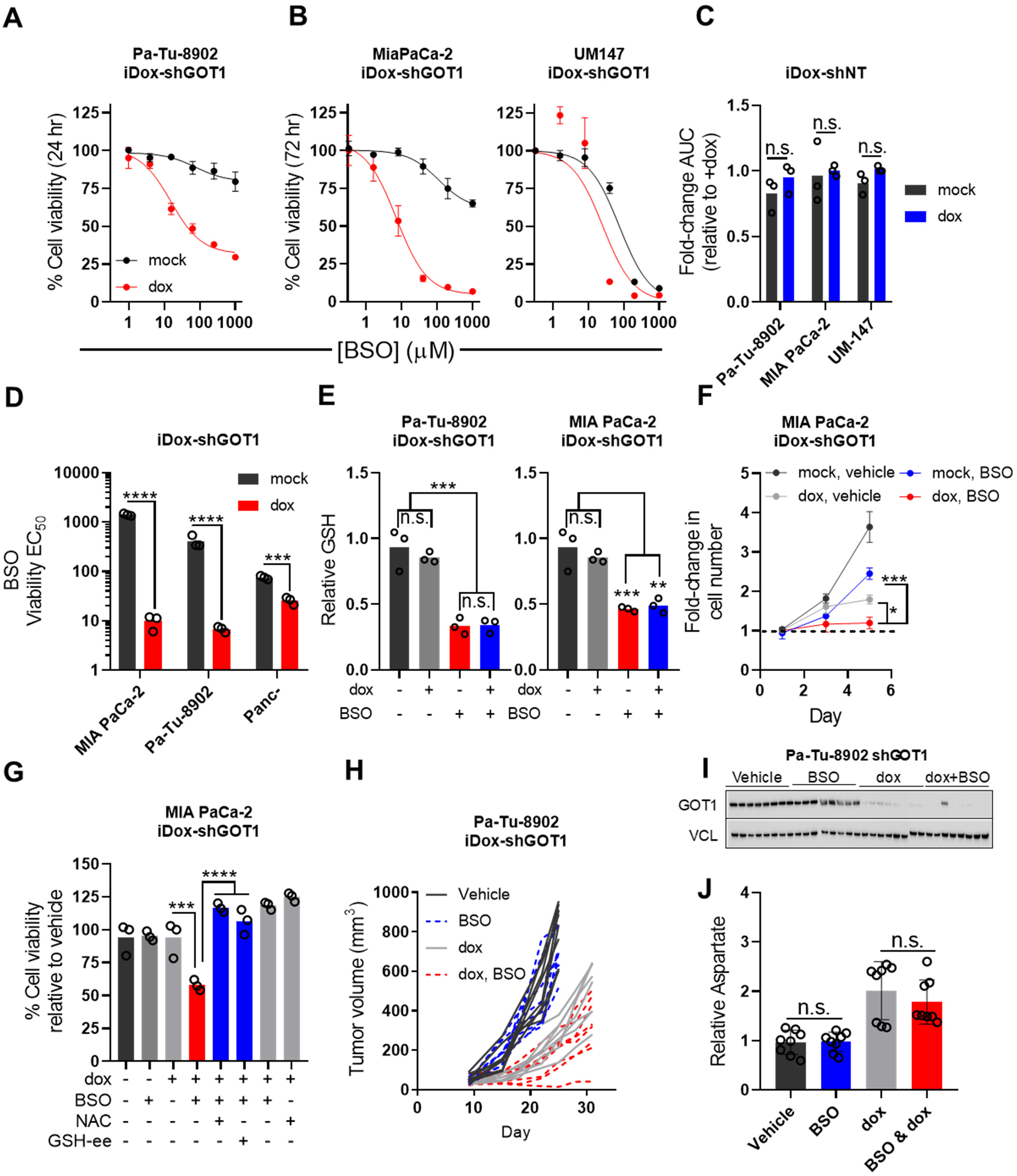
GOT1 knockdown sensitizes PDA to GCLC inhibition. **A-B**) Dose-response curves for iDox-shGOT1 cells after 5 days of GOT1 knockdown followed by BSO treatment for 24 (**A**) or 72 hours across 2 PDA cell lines (**B**) (n=3). **C**) Area under the curve (AUC) normalized to +dox in matched sh non-targeting (shNT). Cells were cultured with dox for 5 days prior to drug treatment. Viability is normalized to a 0.1% DMSO vehicle control (n=3). **D**) Raw EC_50_ values corresponding to Figure 3B and Figure S3B (n=3). **E**) Relative GSH levels normalized to viability and vehicle after 6 hours of 40μM BSO treatment (n=3). **F**) Fold change of MIA PaCa-2 iDox-shGOT1 cell numbers after 5 days of GOT1 knockdown and treatment with the indicated conditions. 40μM of BSO was administered on day 1. Cell numbers are normalized to day 1 for each condition (n=3). **G**) Relative viability of MIA PaCA-2 iDox-shGOT1 cells after 5 days of GOT1 knockdown and treatment with 40μM BSO or co-treatment with 0.5mM N-acetyl cysteine (NAC) 0.5mM GSH-ethyl ester (GSH-EE, n=3). **H**) Individual tumor volume measurements corresponding to Figure 3F (n=8). **I**) Immunoblot analysis of GOT1 from tumors in Figure 3F (n=8). **J**) Relative abundances of aspartate from whole tumor metabolite extracts from tumors in **Figure 3E** (n=8). Error bars represent mean ± SD. Two-tailed unpaired T-test or 1-way ANOVA: Non-significant P > 0.05 (n.s. or # as noted), P ≤ 0.05 (*), ≤ 0.01 (**), ≤ 0.001 (***), ≤0.0001 (****). See also **Figure 3**.

**Supplementary Figure 4, related to Figure 4.**
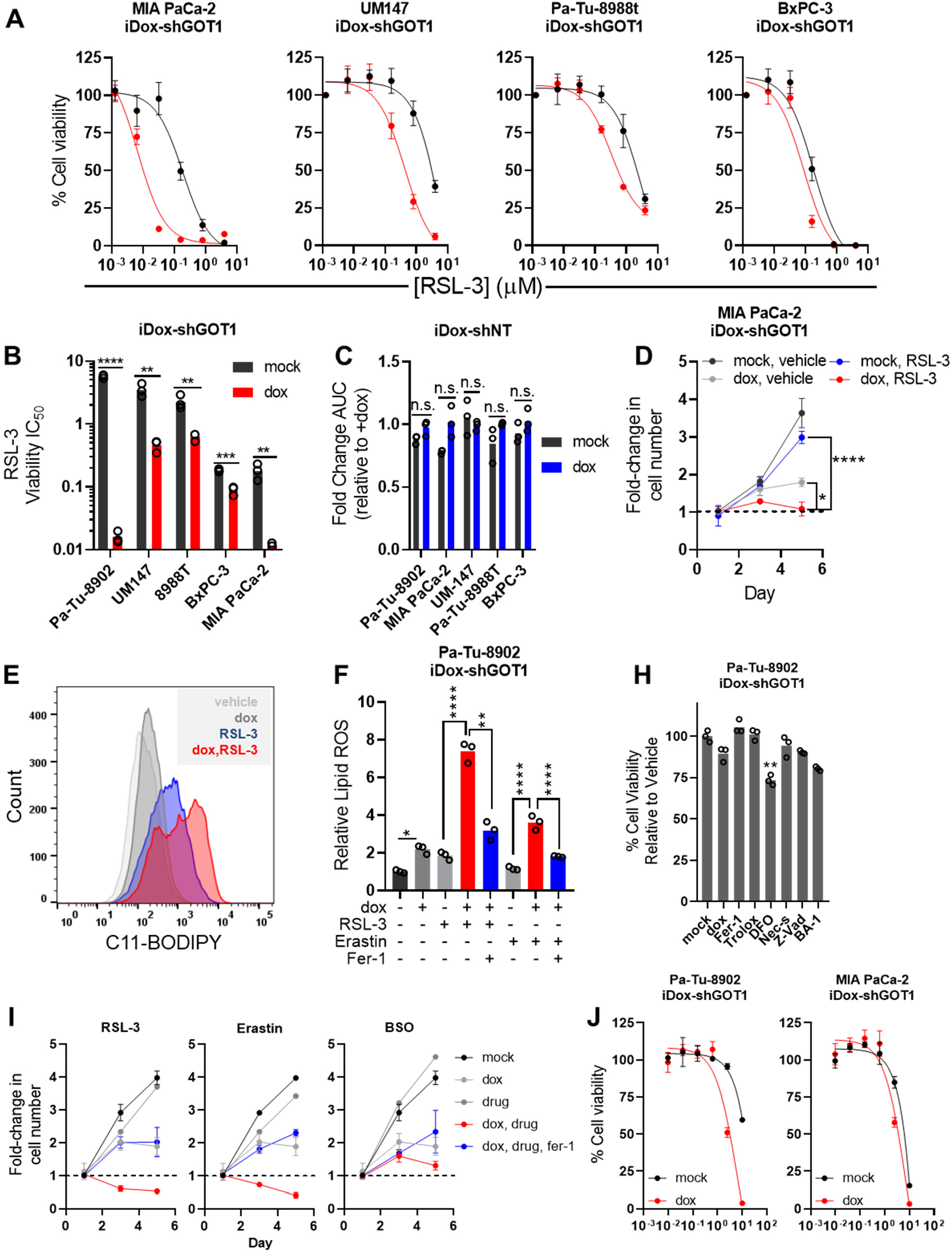
GOT1 suppression sensitizes PDA to ferroptotic cell death. **A**) Percent cell viability dose-response curves from 5 additional shGOT1 PDA cell lines upon 24 hours of RSL-3 treatment. GOT1 was knocked down 5 days prior to drug treatment. Viability is normalized to a 0.1% DMSO vehicle control (n=3). **B**) Raw EC_50_ values for cells treated with RSL-3 for 24 hours. **C**) Area under the curve (AUC) normalized to +dox in matched sh non-targeting (shNT) cell lines. Cells were cultured with dox for 5 days prior to drug treatment. Viability is normalized to a 0.1% DMSO vehicle control (n=3). **D**) Fold change Mia PaCa-2 iDox-shGOT1 cell numbers following 5 days of knockdown and treatment with the indicated conditions. 32nM of RSL-3 was administered on day 1. Cell numbers are normalized to day 1 for each condition (n=3). **E**) Distribution of Pa-Tu-8902 iDox-shGOT1 cells positive for C11-BODIPY corresponding to Figure 4E. Data are normalized to the -dox and vehicle-treated condition (n=3). **F**) Relative lipid ROS in Pa-Tu-8902 iDox-shGOT1 treated with 32nM RSL-3 or 750nM Erastin -/+ 1μM Ferrostatin-1 (Fer-1) for 6 hours (n=3). **H**) Single agent cell viability controls corresponding to **Figures 5F** and **SFigure 5G**. Data are normalized to the -dox and vehicle treated control (n=3). **I**) Fold change Pa-Tu-8902 iDox-shGOT1 cell numbers following 5 days of knockdown and treatment with the indicated conditions (n=3). 32nM RSL-3, 750nM Erastin, or 40μM BSO -/+ 1μM Ferrostatin-1 (Fer-1) were used. Cell numbers are normalized to day 1 for each condition (n=3). **J**) Cell viability dose-response curves upon 24 hours of (–)–FINO_2_ treatment following 5 days of GOT1 knockdown (n=3). Viability is normalized to a 0.1% DMSO vehicle control. Error bars represent mean ± SD. Two-tailed unpaired T-test or 1-way ANOVA: Non-significant P > 0.05 (n.s. or # as noted), P ≤ 0.05 (*), ≤ 0.01 (**), ≤ 0.001 (***), ≤0.0001 (****). See also **Figure 4**.

**Supplementary Figure 5, related to Figure 5.**
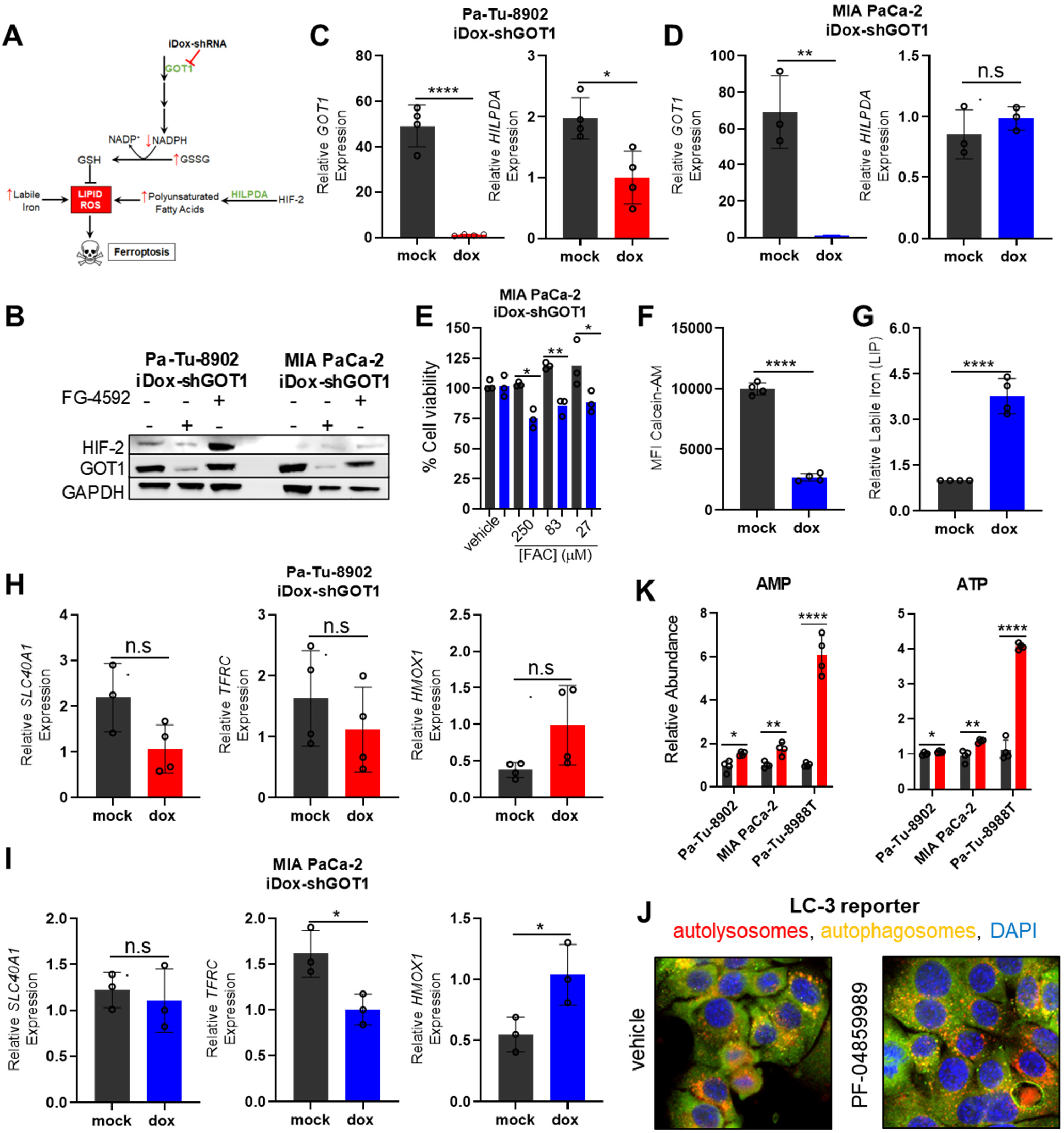
GOT1 knockdown promotes labile iron release in response to metabolic stress. **A**) Model depicting known metabolic regulators of ferroptosis. **B**) Immunoblot for HIF-2 in response to GOT1 knockdown. FG-4592 is a positive control for HIF-2 expression. **C-D**) Gene expression (mRNA) of HILPDA in Pa-Tu-8902 (**C**) and MIA PaCa-2 (**D**) following knockdown (n=4 in C, and n=3 in D). **E**) Cell viability dose-response curves upon 72 hours of ferrous ammonium citrate after 5 days of GOT1 knockdown (n=3). Viability is normalized to a 0.1% DMSO vehicle control. **F-G**) Mean fluorescence intensity (MFI) (**F**) Calcein-AM in MIA PaCa-2 iDox-shGOT1 and relative labile iron (**G**) (n=4). **H-I**) mRNA expression of iron regulatory genes in Pa-Tu-8902 (**H**) and MIA PaCa-2 (I) iDox-shGOT1 following 5 days of knockdown (n=3). **J**) Representative images of basal autophagic flux in MT3-LC3 tandem fluorescence (GFP–RFP) reporter treated with GOT1 inhibitor PF-04859989. **K**) AMP and ATP levels corresponding to **Figure 5H** measured by liquid-chromatography tandem mass spectrometry, normalized to -dox (n=4). Error bars represent mean ± SD. Two-tailed unpaired T-test or 1-way ANOVA: Non-significant P > 0.05 (n.s. or # as noted), P ≤ 0.05 (*), ≤ 0.01 (**), ≤ 0.001 (***), ≤0.0001 (****). See also **Figure 5**.

**Supplementary Figure 6.**
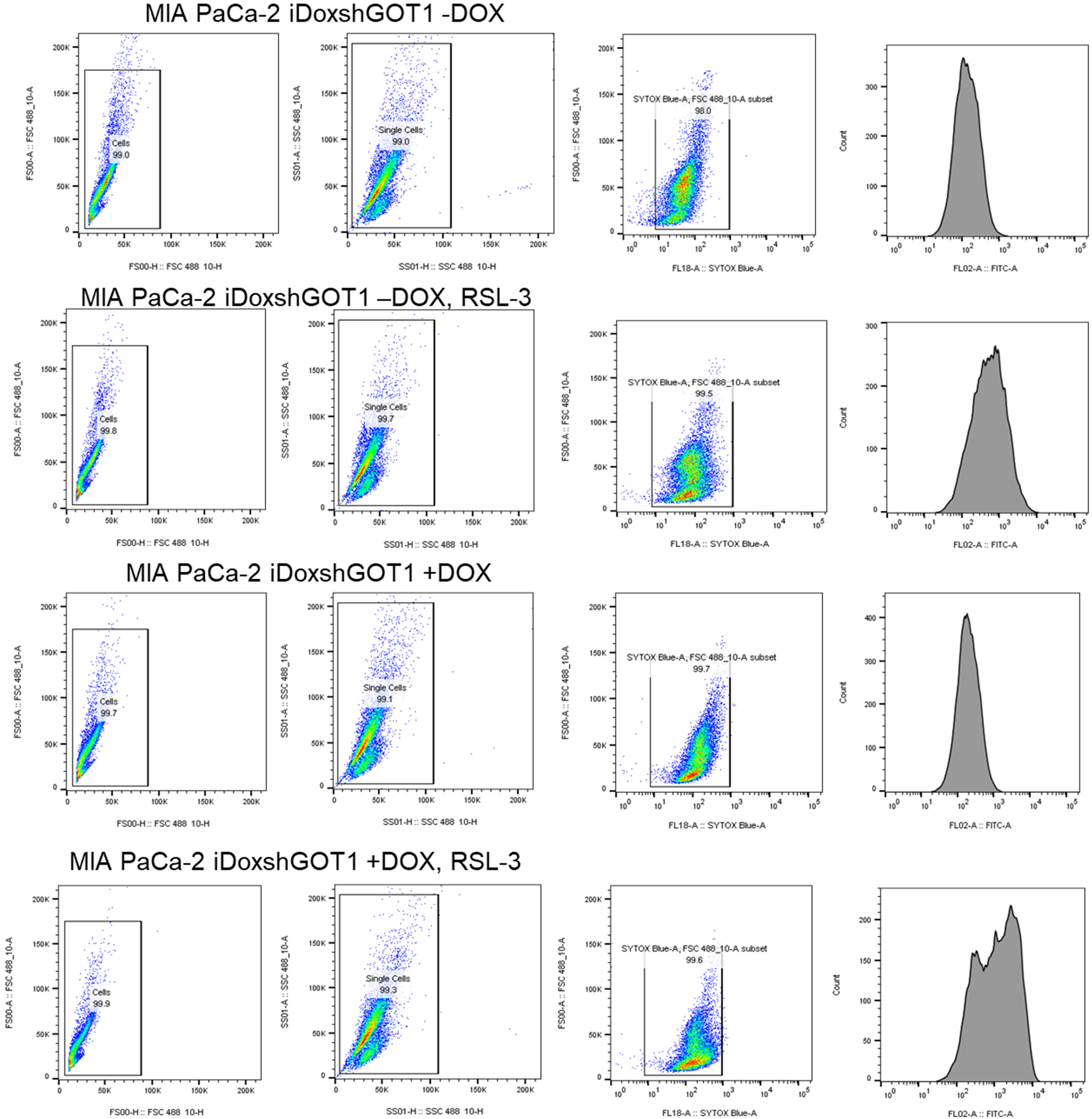
Representative C11-BODIPY Gating. Cells were first gated by size using forward scatter area (FSA) vs. forward scatter height (FSH). Doublet exclusion was then performed using side scatter area (SSA) vs. side scatter height (SSH). Live cells were assessed selecting the Sytox Blue (DAPI) negative population in both experiments. C-11 BODIPY (FITC) were measured in single and live cells for each condition above.

**Supplementary Figure 7.**
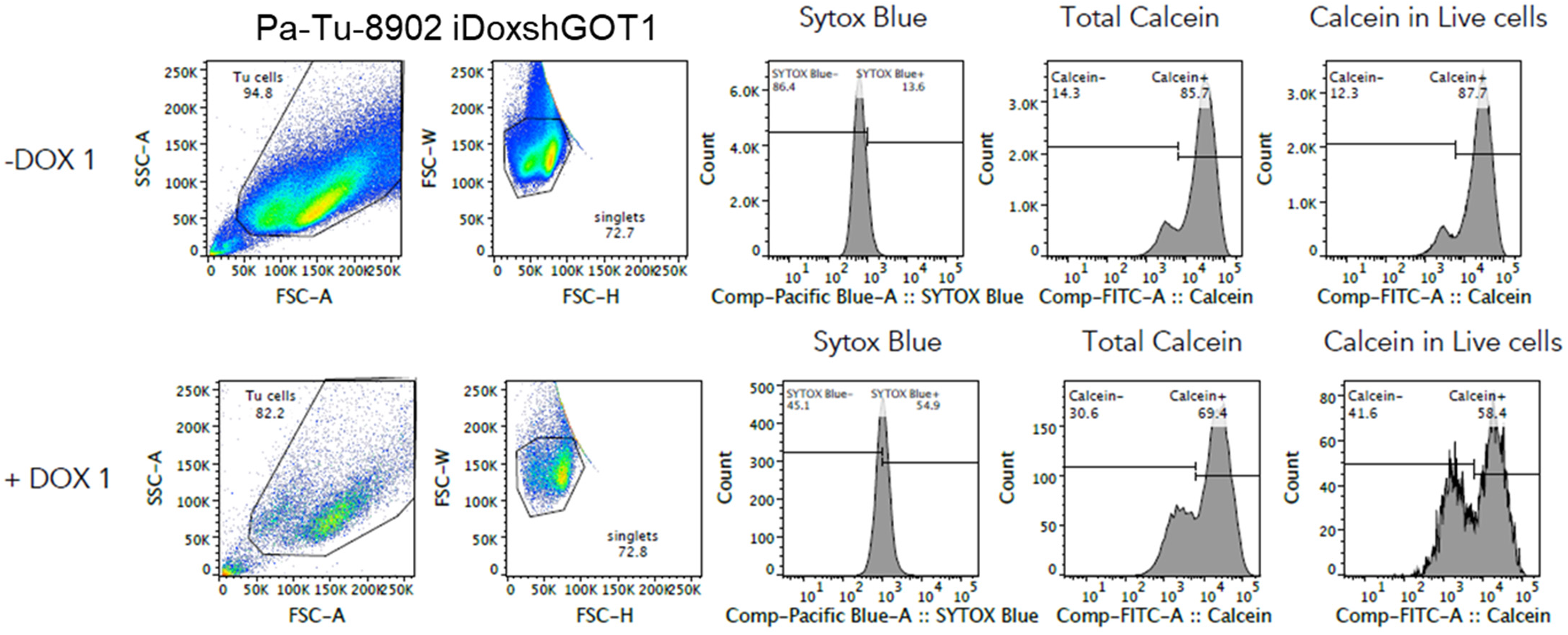
Representative Calcein-AM Gating. Cells were first gated by size using side scatter area (SSC-A) vs. forward scatter area (FSC-A). Doublets were excluded by comparing forward scatter width (FSC-W) vs. forward scatter height (FSC-H). Live cells were assessed selecting the Sytox Blue (Pacific Blue) negative population in both experiments. Calcein-AM (FITC) levels were measured in single and live cells.

